# Generation and maintenance of apical rib-like actin fibers in epithelial support cells of the *Drosophila* eye

**DOI:** 10.1101/2025.05.09.653141

**Authors:** Abhi Bhattarai, Emily W McGhie, Joshua C Woo, Srijana Niraula, Patrick Rosetti, Jaxon M Kim, Ezekiel Popoola, Ruth I Johnson

## Abstract

Heterogeneity and complexity of cytoskeletal structures, and how these are regulated, is poorly understood. Here we use cells of the *Drosophila* pupal eye as models to explore diversity in the actin cytoskeleton. We found that different F-actin structures emerge in primary (1°), secondary (2°), and tertiary (3°) pigment cells as they mature. 1° cells became characterized by dense accumulations of apical F-actin that we termed Apical Ribs of Actin Fibers (ARAFs). The formins Diaphanous and Dishevelled Associated Activator of Morphogenesis are essential for generation of ARAFs, which are connected into a network by α-Actinin, the villin Quail, and Spectrins, and linked to the apical membrane by Quail and Spectrins. ARAFs are contractile, stress-fiber-like, and connect to adherens junctions. Impairing ARAFs indicated that this network maintains cortical tension and is crucial for 1°s to achieve their characteristic shapes. Our evaluation of the three-dimensional shape of 1°s reveals that ARAFs are essential for the rounding and elevation of the apical membrane. Hence, a toolkit of conserved actin regulatory proteins builds and maintains a network of apical stress fibers that governs the morphology of the cell.

**Summary Statement:** This study describes a novel F-actin network that shapes the apical region of cells of the *Drosophila* eye and uncovers a set of proteins that build and maintain the network.

## Introduction

Many tissues are characterized by collections of cells with cell-type specific three-dimensional (3-D) architectures. The organ of Corti of the mammalian cochlea, with its distinctly shaped hair cells and associated support cells, is one remarkable example (Lim, 1986). The mammalian intestine and airway epithelia similarly contain cells that can be easily identified because of their distinct morphologies (Davis and Wypych, 2021; Eenjes et al., 2022; Hewitt and Lloyd, 2021). The *Drosophila* eye, in which the different cell types acquire characteristic shapes during pupal development, is an invertebrate example and an experimental model in which mechanisms that determine cell-specific architectures can be interrogated.

The cells of the *Drosophila* compound eye emerge from a common lineage during larval and pupal development so that each cell type can be unambiguously identified by its morphology and position by 30 - 40 hour after puparium formation (APF) (Cagan and Ready, 1989; Ready et al., 1976; Tomlinson and Ready, 1987; Waddington and Perry, 1960; reviewed by Charlton-Perkins et al., 2021; Johnson, 2021; Kumar, 2012; Pichaud, 2014; Treisman, 2013). Eight photoreceptors form the core of each ommatidium of the eye, surrounded by four epithelial cone cells (CCs; also known as Semper’s cells) and two primary pigment cells (1°s), and a lattice of secondary (2°) and tertiary (3°) pigment cells and mechanosensory bristles separates the ommatidia (Fig. 1A,B). When adherens junctions (AJs) between cells are detected, stereotypical cell shapes are observed as follows at 40 h APF: two fabiform 1°s connect to form a ring around the CCs of each ommatidium; the CCs are rounded and organized with dorsal and ventral CCs in contact with one another; 2° cells are rectangular and those placed horizontally between ommatidia are wider than those positioned obliquely; 3° cells are placed at vertices between three neighboring ommatidia and are hexagonal.

**Figure 1:**
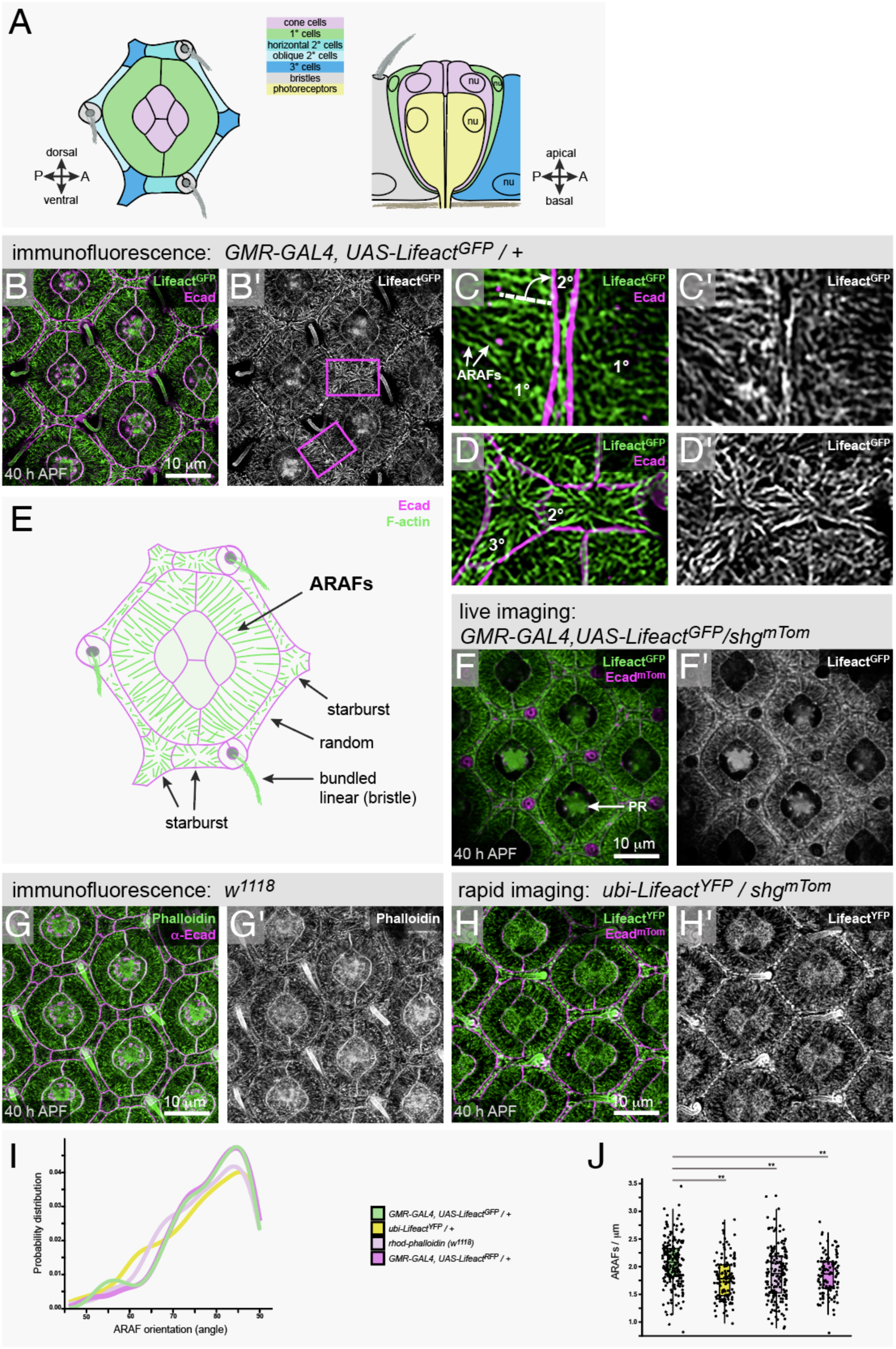
Cell-type specific actin structures are observed in epithelial support cells of the *Drosophila* pupal eye. (A) Cartoons of the apical face (left) of a single ommatidium at 40 h APF and (right) the ommatidium sliced longitudinally, with nuclei (nu) indicated. (B) Small region of an eye, with *Lifeact^GFP^* driven by *GMR-GAL4*. Ecad detected in magenta. Boxed regions in B’ are presented at higher magnification in (C) and (D). In (C), white arrows indicate two almost-parallel ARAFs, white dotted line drawn at 79° to AJ indicates mean orientation of ARAFs with respect to the 1°:LC border. (E) Cartoon summarizing apical-cortical F-actin organization in an ommatidium. Cartoon excludes actin in CCs and at ZAs. (F) Small region of the eye of a live pupa, with *Lifeact^GFP^* and endogenous Ecad^mTom^ (encoded in *shgm^Tom^*). Bristles were vertical and hence not captured in this image. *Lifeact^GFP^*was not driven in all CCs. Lifeact^GFP^-rich photoreceptor rhabdomeres (PR) indicated with white arrow. G) Ommatidia of a *w^1118^*eye, with F-actin detected with phalloidin. (H) Region of an eye with *ubi-Lifeact^YPF^*and Ecad^mTom^. The *ubi* promoter generated higher levels of Lifeact^YFP^ in anterior and posterior CCs. (I) Distribution plot of the angle at which ARAFs oriented toward the 1°:LC boundary. (J) Analyses of the density of ARAFs in 1° cells. Asterisks indicate significant differences (see Tables S1and S2). All scale bars are 10 μm.

As with any cell, the shapes that *Drosophila* eye cells acquire are governed by a multitude of factors. These include the shape, density and stability of the cytoskeleton, actomyosin activity, composition and stability of junctions, placement of organelles, and forces transmitted to the cell from neighboring cells and the extracellular environment. Only a subset of these factors have been studied in the *Drosophila* eye (Johnson, 2021). In particular, how the cytoskeleton is organized in differently shaped *Drosophila* eye cells has not been examined.

Strategies to visualize the actin cytoskeleton rely on introducing fluorescent molecules into the cell that bind actin filaments (F-actin). Tagged phalloidin remains the benchmark reagent for fixed preparations (Wulf et al., 1979), but for imaging F-actin in live tissues the standard strategy is to introduce fusion proteins into the tissue composed of actin binding domains (ABDs) linked to fluorescent proteins. ABDs used include the 280 amino acid tail of *Drosophila* Moesin (Edwards et al., 1997), the 255 amino acid calponin homology domain region of human Utrophin (Burkel et al., 2007), a 43 amino acid peptide from rat inositol 1,4,5-trisphosphate 3-kinase A termed F-tractin (Johnson and Schell, 2009), and a 17 amino acid peptide of yeast Abp140 termed Lifeact (Riedl et al., 2008). These ABD fusion proteins have been engineered as expression transgenes in *Drosophila* (Bloor and Kiehart, 2001; Cai et al., 2014; Dutta et al., 2002; Edwards et al., 1997; Hatan et al., 2011; Huelsmann et al., 2013; Kiehart et al., 2000; Spracklen et al., 2014; Zanet et al., 2012) and, alongside phalloidin, used to track cytoskeletal organization and dynamics in a variety of *Drosophila* tissues.

*Drosophila* tissues have proven invaluable in revealing ways in which actin, and actomyosin complexes, are deployed to regulate cell behavior and drive development. Examples include polarized lamellipodia, filipodia, and ‘whip-like’ structures that project from the follicular epithelium enclosing each egg chamber and propel collective migration of the follicular cells (Cetera et al., 2014; Squarr et al., 2016); filopodia that, during dorsal closure, guide fusion of apposing epithelial sheets (Jacinto et al., 2000; Millard and Martin, 2008); cytonemes that project to mediate signal transduction across and between tissues (Casas-Tinto and Portela, 2019). Examples of extraordinary subcellular actin organization include the large arrays of actin cables in nurse cells that position the nuclei (Huelsmann et al., 2013; Logan et al., 2022), and the supracellular actomyosin rings in tracheal cells that ensure the tracheal lumens are correctly formed (Hannezo et al., 2015; Matusek et al., 2006; Ozturk-Colak et al., 2016). Examples of stress fibers include those of follicle cells of the egg chamber (Cetera et al., 2014; Delon and Brown, 2009) that may guide the trajectory of collectively migrating follicle cells (Sherrard et al., 2021), generate compression that elongates the enclosed oocyte (He et al., 2010; Popkova et al., 2020; Qin et al., 2017), and flatten and expand the follicular epithelium as the oocyte grows (Li et al., 2024). Examples of transient, less-patterned cytoskeletal structures include the cortical-medial actomyosin meshworks of the embryonic mesoderm (Martin et al., 2009), epithelium (Fernandez-Gonzalez and Zallen, 2011; Rauzi et al., 2010; Sawyer et al., 2011), amnioserosa (Blanchard et al., 2010; David et al., 2010; Solon et al., 2009), and salivary gland (Booth et al., 2014). Pulsing of these actomyosin meshworks mediates cell and tissue shape changes whilst maintaining tissue integrity (see Blanchard et al., 2018; Coravos et al., 2017; Miao and Blankenship, 2020; Perez-Vale and Peifer, 2020; Sutherland and Lesko, 2020). Finally, stable, polarized, supracellular actomyosin cables associated with cell junctions are observed in various embryonic and larval tissues, and align cells and generate boundaries that separate groups of cells (Fernandez-Gonzalez and Harris, 2023; Harris, 2018; Roper, 2015).

We set out to characterize the actin cytoskeleton in epithelial cells of the *Drosophila* pupal eye with the hypothesis that F-actin would be employed differently to support the stereotypical shapes that each cell type acquires. We found that the apical cortical cytoskeleton was organized into ‘starburst’ patterns in 3°s and horizontal 2° cells, and F-actin was randomly organized in oblique 2°s. In contrast, 1° cells become packed with an organized mass of actin cables that tracked across the cells’ width. We term these cables Apical Ribs of Actin Fibers (ARAFs) and document a set of actin regulators that generate, organize and maintain the ARAFs. Several of these regulators were present at higher levels in 1° cells than in 2°s and 3°s. The ARAFs connect to AJs, and our data indicated that they are tensile and essential for the fabiform apical shape of 1°s. To the best of our knowledge, ARAFs are the third reported instance of apical stress fibers in *Drosophila* epithelia: these have previously been described in tracheal cells and the thoracic epithelium (Hannezo et al., 2015; Lopez-Gay et al., 2020). 3-D analyses of the morphology of 1°s revealed that ARAFs also maintain the elevated dome shape of the ommatidium, observed in longitudinal sections. Taken together, this study provides a conceptual framework for the genetic control of a tensile cytoskeletal network that is essential for a complex cell shape to emerge in an epithelium.

## Results

### Cell-specific apical-cortical actin structures are observed in the *Drosophila* pupal eye

Previous detection of F-actin with phalloidin had suggested that the cytoskeleton becomes increasingly complex in epithelial cells of the *Drosophila* pupal eye as they adopt their characteristic shapes (Johnson et al., 2008). The fly eye has four types of epithelial support cells: 1°, 2° and 3° pigment cells, and CCs (Fig. 1A,B). These cells enshroud the photoreceptor clusters at the core of each ommatidium to provide mechanical support, optically separate each ommatidium, and generate extracellular material that becomes organized into lenses that will cap each ommatidium. We drove expression of *UAS-Lifeact^GFP^* with *GMR-GAL4* (Freeman, 1996; Hatan et al., 2011), and coupled this approach with high-resolution confocal microscopy to resolve in detail how the actin cytoskeleton is organized in the epithelial support cells when they achieve their stereotypical shapes (Fig. 1B-D, summarized in Fig. 1E).

Our approach revealed numerous apical-cortical actin filaments, organized across the width of the fabiform 1° cells (Fig. 1B,C,E) that we name Apical Ribs of Actin Fibers (ARAFs) to describe their morphology. Most ARAFs were oriented toward the boundary between 1° and surrounding 2° or 3° cells at ∼84° (Fig 1I, median = 79.49°, n = 259 ARAFs, analyzed in 20 1° cells). In 3°s, which are hexagonal in outline, the apical-medial F-actin network was organized with filaments tracking toward the approximate cell center in a ‘starburst’ formation (Fig. 1D,E). A similar starburst actin formation was observed in 2° cells positioned horizontally about each ommatidium (Fig. 1D,E), and F-actin was randomly oriented in the narrower oblique 2°s (Fig. 1B,E). The *GMR-GAL4* line we used did not consistently drive *UAS-Lifeact^GFP^* expression in CCs, so we excluded these from our analyses.

Apical actin structures fragmented or collapsed readily during sample preparation, so we developed strategies to prevent this (see Materials and Methods). Further, we were concerned that fixation, immunofluorescence, or *Lifeact* expression would modify cytoskeletal organization, but we observed the same F-actin structures in retinas of live pupae (Fig. 1F), retinas incubated with phalloidin (Fig. 1G), retinas expressing low levels of *Lifeact* driven with the *ubiquitin* promoter (Fig. 1H, Santa-Cruz Mateos et al., 2020) or *sparkling-GAL4* driver (not shown, Jiao et al., 2001), retinas expressing the ABD of Moesin (*UAS-GMA*, data not shown, Bloor and Kiehart, 2001) and retinas fixed and immediately imaged (‘rapid imaging’, Fig. 1H, data not shown). Image resolution was lower when imaging the cytoskeleton in live retinas (Fig. 1F) since the eye is surrounded by hemolymph and still internal to the head with an overlying layer of epithelium through which one must focus. We observed identical actin organization when labelling the cytoskeleton with Lifeact^GFP^ (Fig. 1B,F), Lifeact^YFP^ (Fig. 1H), Lifeact^RFP^ (not shown, Huelsmann et al., 2013), or phalloidin (Fig. 1G), although the actin density differed modestly. For example, ARAFs were identically oriented regardless of detection strategy (Fig. 1I), but more ARAFs were observed in 1°s expressing *Lifeact^GFP^*, fewer with *Lifeact^YFP^*, and ARAF density was most variable when tagged phalloidin was utilized (Fig. 1J). These minor disparities could be due to differences in emission intensity or photostability of the fluorophores: Lifeact^YFP^ rapidly photobleached whilst Lifeact^GFP^ robustly labelled ARAFs; rhodamine-phalloidin and Lifeact^RFP^, which have similar excitation/emission spectra, labelled ARAFs similarly, but less brightly than Lifeact^GFP^ (Fig. 1J, Tables S1, S2). Hence, given the similarities in ARAFs observed with phalloidin and Lifeact, we consider it unlikely that Lifeact significantly alters ARAF architecture. In contrast, more zonula adherens-associated F-actin (ZA-actin) was detected with phalloidin than with Lifeact (compare Fig. 1G’ with Fig. 1B’,H’).

### Formation of ARAFs correlates with rounding of the 1° cell:LC border

Given the extraordinary organization of ARAFs in 1° cells, we next explored how these are generated. We first observed nascent ARAFs in the apical-medial domain of 1°s from around 33 h APF (Fig. 2). Before then, a loose, patchy meshwork of apical F-actin was detected (Figs 2A, B, S2). Gradual emergence of short, and then longer, fibers was observed (Movie S1), and most of these were oriented across the width of 1° cells (Fig. 2B, see 33:40 h APF). In addition, ‘whisker-like tufts’ of F-actin were observed in the dorsal and ventral regions of most 1°s (Fig. 2B 33:40 h APF, Fig. 2C). These tufts became less prominent as the organized ARAF network emerged from around 36 h APF (Fig. 2D). Whether ARAFs were generated from reorganization of extant actin filaments or instead generated *de novo*, remains to be investigated. Prior to 33 h APF, the density of apical-cortical actin in 1° cells remained relatively consistent (Fig. S2B) but as ARAFs emerged, their density rapidly increased (Fig. 2E). Accordingly, before ARAF emergence apical-cortical F-actin was more dense in 2° and 3° cells (collectively called lattice cells, LCs) than in 1°s (Fig. 2B,C, Fig. S2A,C) but actin density became roughly equivalent in these cells from 39 h APF (Fig. 2D,F). Of note, scalloping of the boundaries between 1° cells and LCs was pronounced before 33-34 h APF (see 1° outlined in Figs 2B, S2A), and emergence of the network coincided with gradual relief of scalloping and the adoption of the characteristic rounded shape of 1° cells (Fig. 2D), suggesting that ARAFs support the adoption of this shape.

**Figure 2:**
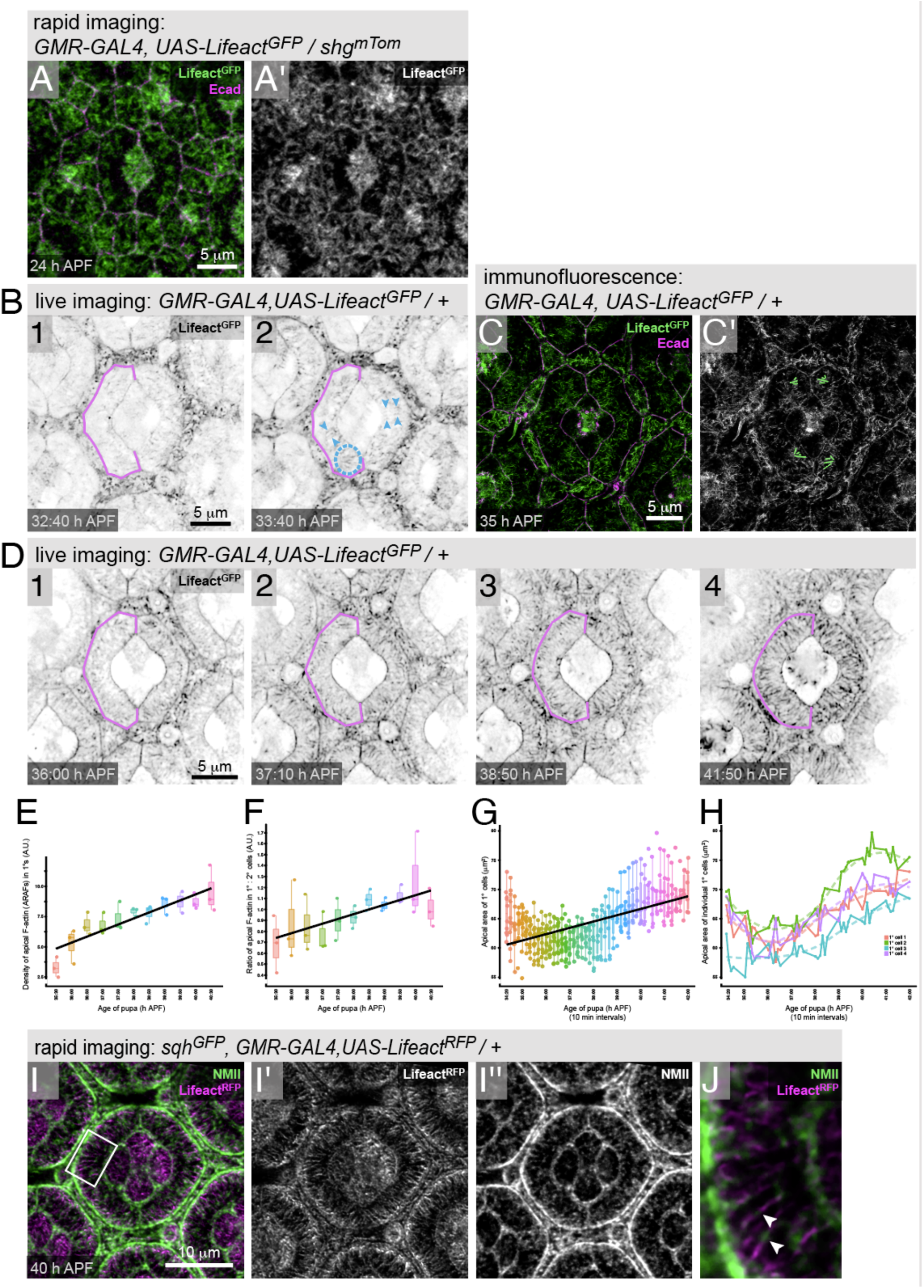
Filaments of the ARAF network emerge from ∼33 h APF. (A) Lifeact^GFP^ and Ecad^mTom^ in an ommatidium at 24 h APF. (B) Two stills (1 and 2) from an eye with Lifeact^GFP^ imaged live, at ages APF as indicated. See Fig. S2 for additional stills. Blue arrowheads indicate groups of emergent ARAFs oriented in parallel. Dotted circle outlines a whisker-like tuft of actin. (C) Lifeact^GFP^ in an eye dissected at 35 h APF, with Ecad detected. Whisker-like tufts of actin traced in green. (D) Four stills from Movie S1 of a live retina with Lifeact^GFP^, at ages as indicated. In (B) and (C) the same 1° is outlined in magenta. Analyses of 1° cells imaged live, as follows: (E) density of apical- cortical F-actin and the emergent ARAF network, (F) ratio of F-actin in 1° and 2° cells, and (G) the apical area of 1° cells. (H) Plot of apical area of four 1° cells, imaged live. (I) Lifeact^RFP^ and Sqh^GFP^ (a proxy for NMII) in an ommatidium at 40 h APF. Boxed region in (I) is presented at higher magnification in (J), with white arrowheads indicating NMII along ARAFs. Scale bars are 5 μm or 10 μm.

The apical face of 1° cells spread to partially cover CCs with an ‘awning’, as illustrated in Figure 1A. Analyses of the area occupied by this apical 1°cell awning (represented as the cross-sectional area of a 1° cell, as imaged at AJs) showed an increase in area as the ARAF network assembled (Fig. 2G). In addition, area of the 1° cell awning oscillated (Fig. 2H). Similar fluctuations in the apical area of 1°s have been reported in younger retinas before ARAF emergence and ascribed to oscillatory assembly of cortical actomyosin meshworks (Blackie et al., 2020). Indeed, we also detected non-muscle myosin II (NMII) within the ARAF network (Fig. 2I), suggesting that the oscillations in size of the 1° cell awnings that we observed are driven by actomyosin activity within the ARAF network, and indicating that this is a tensile network. NMII also accumulated at high levels toward the 1°:LC borders.

### Formation or maintenance of ARAFs is dependent on Formins and Enabled

We predicted that formins were instrumental in generating ARAFs and our survey of the *Drosophila* formins Cappuccino (Capu) (Emmons et al., 1995; Quinlan et al., 2005; Rosales-Nieves et al., 2006), Diaphanous (Dia) (Afshar et al., 2000; Castrillon and Wasserman, 1994; Grosshans et al., 2005), Dishevelled Associated Activator of Morphogenesis (DAAM) (Matusek et al., 2006; Matusek et al., 2008), Formin 3 (Form3) (Tanaka et al., 2004), Formin homology 2 domain containing (Fhos) (Anhezini et al., 2012; Lammel et al., 2014), and Formin-like (Frl) (Bai et al., 2011; Rohn et al., 2011), revealed that DAAM and Dia were present in the pupal eye before and during ARAF generation (Figs 3A-H, S3, data not shown). DAAM and Dia family formins are activated by Rho GTPases (Kuhn and Geyer, 2014), and, consistent with their being activated in 1° cells, a Rho/Rac activity sensor localized to the cortical region of 1°s (as well as LCs) during ARAF assembly (Fig. S3E, Abreu-Blanco et al., 2014).

**Figure 3:**
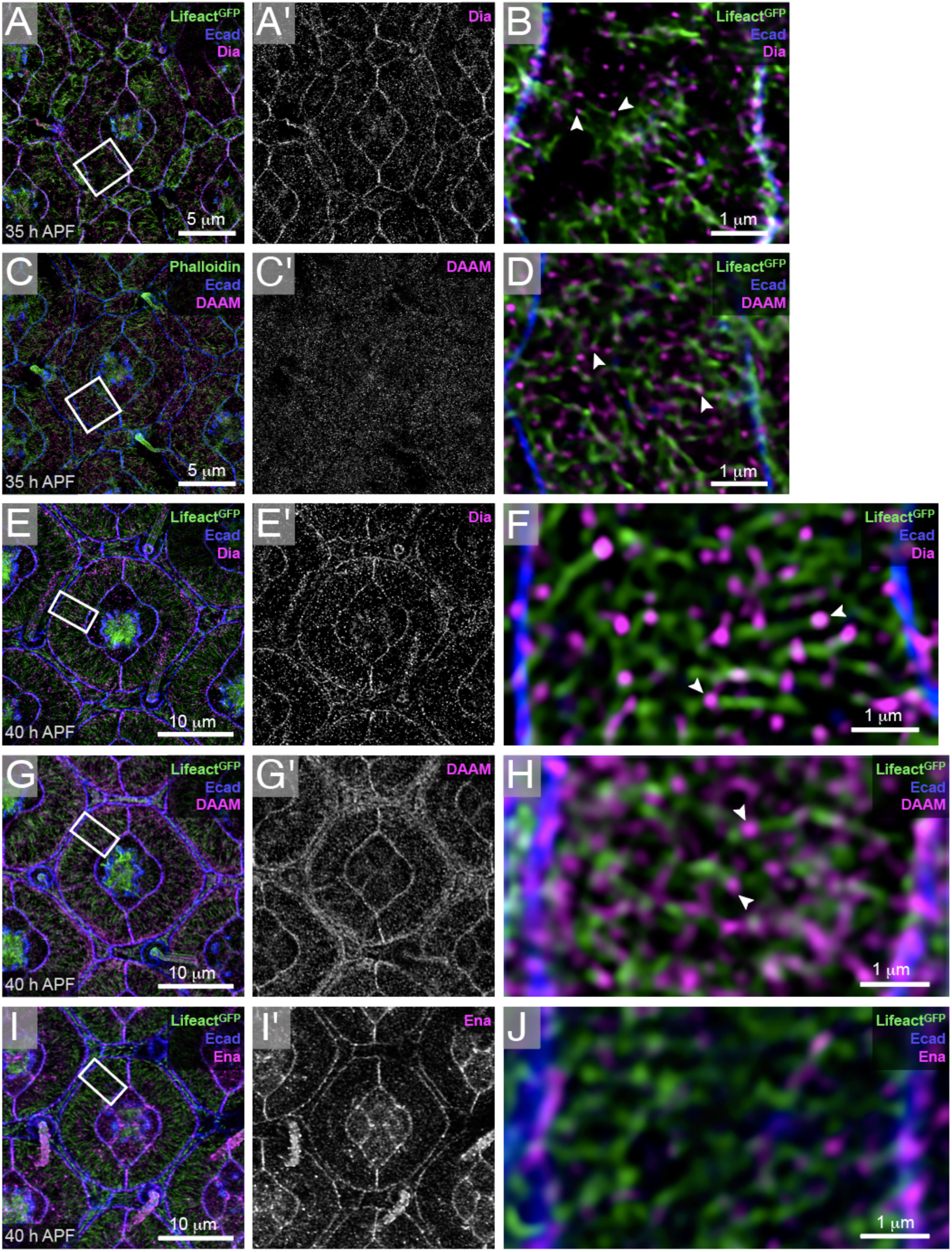
DAAM, Dia and Ena localize to adherens junctions and/or ARAFs. (A) Dia in an ommatidium at 35 h APF, with boxed region at higher magnification in (B). (C) DAAM in a 35 h APF ommatidia, with boxed region in (D). (E) Dia and (G) DAAM in ommatidia at 40 h APF, with boxed regions in (F) and (H) at high magnification, respectively. White arrowheads in (B), (D), (F) and (H) point to Dia or DAAM detected at the tips of ARAFs. (I) Ena in an ommatidium at 40 h APF, with boxed region at higher magnification in (J). Scale bars are 1 μm, 5 μm or 10 μm.

Formins attach to and extend the barbed ends of F-actin (Breitsprecher and Goode, 2013; Courtemanche, 2018), and since Dia and DAAM localized to many short actin fibers oriented across the 1° cell width at 35 h APF (Fig. 3B,D) and 40 h APF (Fig. 3F,H), we predict that the ARAF network is composed of numerous actin cables that tile across 1°s, rather cables that traverse the entire 1° cell width. Dia also accumulated at AJs between 1° cells and their neighbors (Fig. 3A,E), consistent with the role of Dia family formins in supporting ZA-actin and AJ stability (Carramusa et al., 2007; Homem and Peifer, 2008; Sahai and Marshall, 2002). In contrast, DAAM accumulated at AJs by 40 h APF but was not observed at junctions at 35 h APF (Fig. 3D,G). Another barbed-end actin polymerizer, Enabled (Ena, the *Drosophila* vasodilator-stimulated phosphoprotein) (Gertler et al., 1995; Winkelman et al., 2014), localized to AJs throughout the retina (Fig. 3I,J), consistent with AJ-localization in other tissues and cultured cells (Faix and Rottner, 2022).

To test the contributions of Dia and DAAM to ARAF formation, we next reduced their translation using RNA-interference (RNAi). In our hands, when RNAi transgenes are driven with *GMR-GAL4*, protein expression is seldom impacted during larval or early pupal development. Hence specification and earlier morphogenesis of photoreceptors and epithelial support cells are unimpeded. However, by the time of ARAF formation, protein levels are significantly reduced (Table S3). Reducing Dia (by 83.3% by 40 h APF, Table S3) severely disrupted the ARAF network: only sparse fragments of disorganized F-actin were observed in 1° cells by 40 h APF (Fig. 4B,E, Table S4). *DAAM^RNAi^* transgenes less efficiently reduced DAAM (by 39.7%, Table S3), but ARAFs were nonetheless shorter, less dense, and randomly oriented (Fig. 4C,E, Table S4).

**Figure 4:**
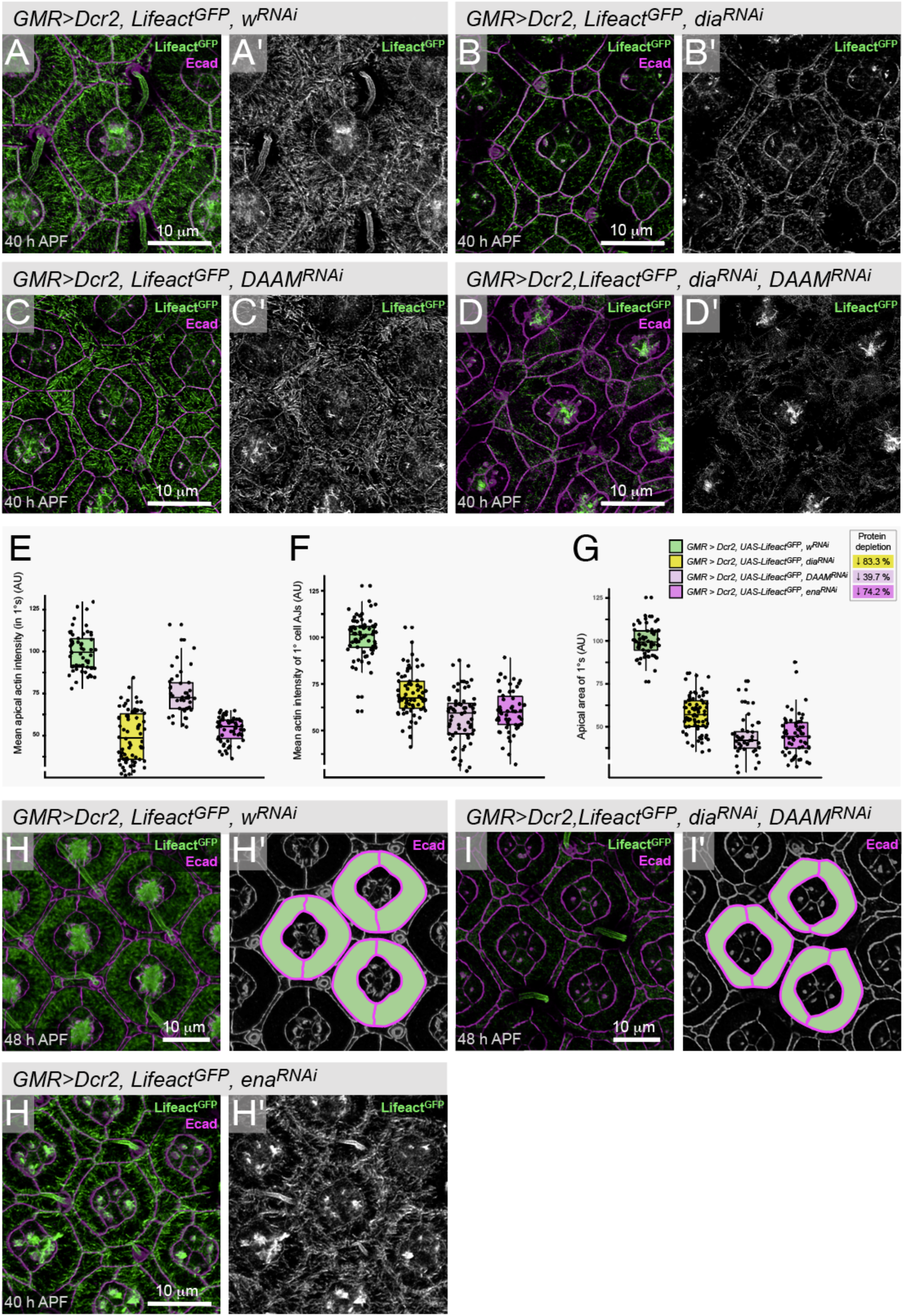
ARAFs are generated and maintained by Formins. (A) Ommatidium in a control retina with *w^RNAi^* transgene expression. F-actin was detected via co-expression of *Lifeact^GFP^*. Ecad labels AJs (magenta). Ommatidia from eyes with (B) *dia^RNAi^*, (C) *DAAM^RNAi^*, or (D) both *dia^RNAi^* and *DAAM^RNAi^*. Analyses of (E) the intensity of Lifeact^GFP^ in the apical awnings of 1°s, a proxy for F-actin/ARAF density, (F) Lifeact^GFP^ detected at ZA-actin at junctions between 1° cells and neighboring 2° and 3° cells and (G) the apical transverse area occupied by the 1° cells, normalized to 100 μm^2^. Statistical test scores relating to the analyses summarized in (E) to (G) are presented in Tables S4 to S6. (H) Ommatidia in a control eye and (I) an eye with *dia^RNAi^* and *DAAM^RNAi^* expression, at 48 h APF. 1° cells are traced in (H’) and (I’) to highlight the cells’ shapes. (J) Ommatidia in an eye with *ena^RNAi^* expression. Scale bars are 10 μm.

Reducing both Dia and DAAM simultaneously obliterated the ARAF network (Fig. 4D).

Reducing these formins also affected ZA-associated actin, which was severely reduced in retinas expressing both *dia^RNAi^*and *DAAM^RNAi^* transgenes, and, surprisingly, more significantly reduced in retinas with reduced DAAM than those with reduced Dia (Fig. 4B’,C’,D’,F and Table S5). Hence, despite being absent from AJs before 40 h APF, DAAM has a more prominent role than Dia in generating ZA-actin in 1° cells of the pupal eye. Indeed, many 1° cell pairs failed to fully encircle the CCs and securely adhere to each other (Fig. 4C), suggesting that DAAM is particularly crucial to secure adhesion between 1°s.

1° cells were severely misshapen when *dia* or *DAAM* expression was reduced: instead of adopting stereotypical rounded outlines, boundaries between 1° and LCs remained straight or scalloped (Fig. 4B-D), and the apical area of 1° cells was consistently smaller (Fig. 4G, Table S6). A caveat with these observations is that RNAi transgenes were expressed across the eye, also disrupting LCs which failed to narrow and consequently non-autonomously affected 1° cell morphology. To address whether misshapen 1° cells were merely developmentally delayed, we next examined the 1°s in older retinas (Fig. 4H,I). Indeed, 1°s did become rounded by 48 h APF, but their width was less regular, and the fabiform shape distorted, even when LCs had narrowed and organized into a hexagonal lattice (Fig. 4I).

Despite not localizing to ARAFs, reducing Ena (by 74.2%, Table S3) also disrupted the ARAF network: fewer ARAFs were present but those that remained were long (Fig. 4H, Table S4). However, ZA-actin was severely reduced in 1°s (Fig. 4F,H, Table S5) and this is consistent with a previously-reported role for Ena/VASP family proteins in assembly or maintenance of ZA-actin (Baum and Perrimon, 2001; Leerberg et al., 2014; Scott et al., 2006; Yu-Kemp et al., 2017). Hence, we posit that Ena does not contribute to ARAF elongation but instead to the generation of ZA-actin that is required for the formation or maintenance of a dense ARAF network. This implies that the ARAF network interacts with ZA-actin.

### The ARAF network connects to the zonula adherens

To test our hypothesis that ARAFs synapse with the ZA in 1°s, we directly disabled ZAs by reducing Ecadherin (Ecad, encoded by *shotgun* in *Drosophila*, the backbone of the AJ) or α-Catenin (α-Cat, which links Ecad to F-actin to stabilize AJs) (Mege and Ishiyama, 2017). Expression of RNAi transgenes generated 1° cells with sparse AJs between 1°s and their 2° or 3° cell neighbors (Fig. 5B,C). Because *GMR-GAL4* drove low (or no) RNAi expression in CCs, AJs at 1°:CC boundaries were less disrupted. Reducing α-Cat in addition generated numerous cytoplasmic puncta of Ecad that we predict are endocytic vesicles containing Ecad that failed to be stabilized in AJs in the absence of abundant α-Cat (Fig. 5C).

**Figure 5:**
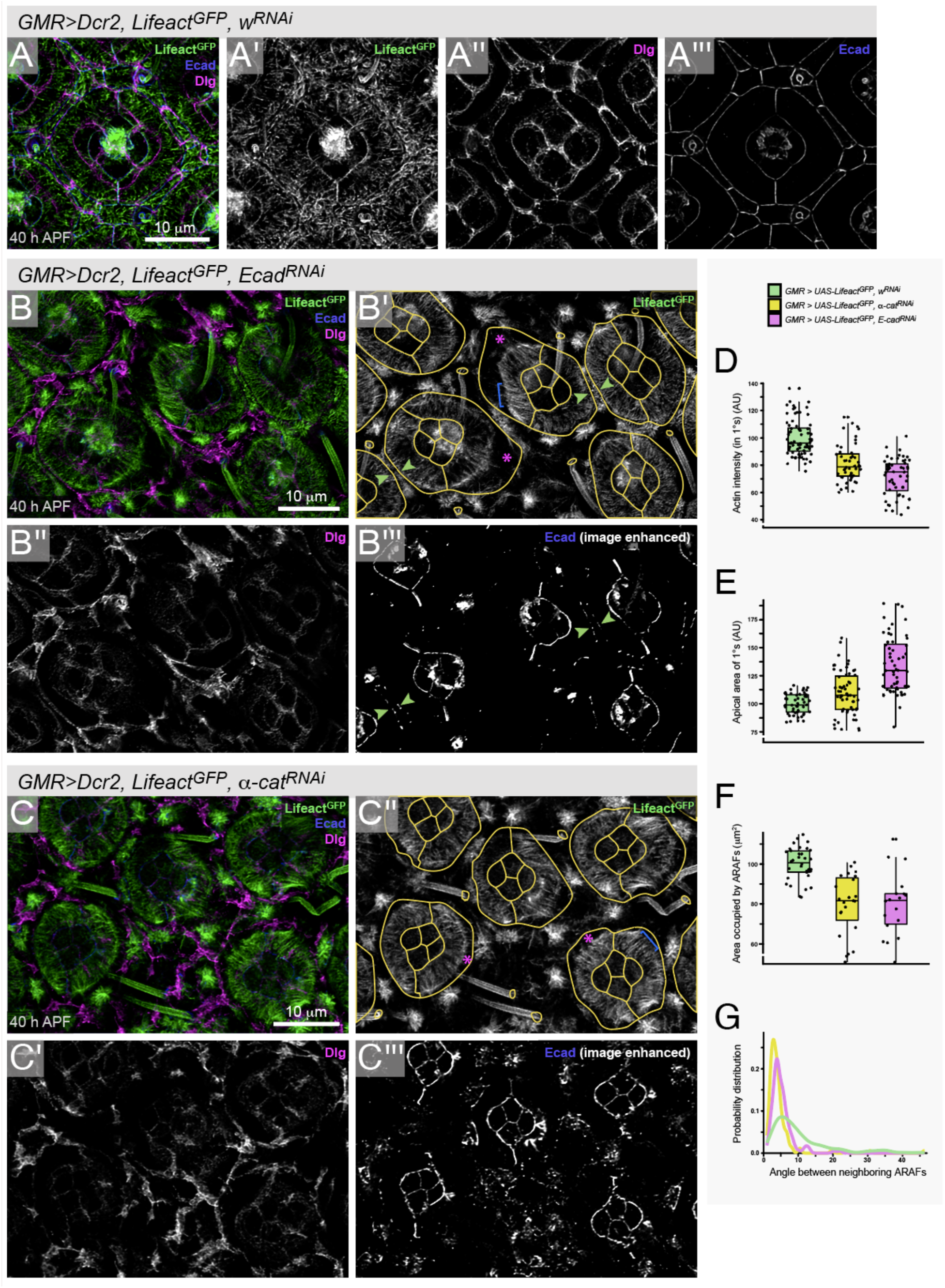
The ARAF network connects to the zonula adherens. (A) Control ommatidium with *w^RNAi^* expression, with (A’) Lifeact^GFP^, (A’’) Dlg in magenta and (A’’’) Ecad in blue. (B) Small region of an eye with with *Ecad^RNAi^* or (C) *α-cat^RNAi^*. For (B’) and (C’), ommatidia, and the bristle collars, are traced in yellow in the Lifeact^GFP^ images, and magenta asterisks highlight examples where the 1° cell membrane bulges away from the edge of the ARAF network, which was released from the membrane. Blue bracket indicates examples of almost-parallel ARAFs. For in (B’’’) and (C’’’) Ecad signal was artificially enhanced to make residual Ecad visible. Green arrowheads in (B’’’) indicate where sufficient residual AJs remain so that ARAFs still connect to the 1° cell AJ. Analyses of (D) the intensity of apical actin (ARAFs) in 1° cells, (E) the area occupied by the 1°s or (F) the ARAF network, and (G) distribution of the orientation of neighboring ARAFs. Statistical test scores relating to (D) to (G) are presented in Tables S7 to S10. Scale bars are 10 μm.

In 1° cells of these retinas, the ARAF network was modified in profound ways (Fig. 5B- D, Table S7). First, whilst ARAFs remained connected to the inner 1° cell boundaries abutting CCs where AJs were retained, large gaps emerged between the ARAF network and the 1°:LC boundaries wherever AJs were especially sparse or absent (Fig. 5B,C, asterisks). 1° cells bulged into the surrounding tissue at these gaps, distorting their shape and enlarging their apical area (Fig. 5E, Table S8). In 1°s where the ARAF network was entirely disconnected along the length of the outer 1° cell border, the network was shaped into an almost-perfect half-ring that echoed the rounded shape of mature 1°s (Fig. 5B,C, asterisks), but occupied a smaller area (Fig. 5F, Table S9), supporting our assertion that ARAFs are contractile, yet stretched across the width of 1° cells. Hence, ARAFs constrain the apical awning of 1°s into their characteristic rounded shape. Remarkably, the presence of just a few AJs along the outer 1° boundary was sufficient to tether the ARAF network and constrain the 1° cell shape (Fig. 5B, arrowheads).

Second, disconnecting ARAFs from AJs altered organization of the network. In control retinas, most ARAFs were oriented at angles of ∼6° between neighboring fibers so that they ‘fanned’ toward the outer 1°:LC boundary (Fig. 5A,G). Disrupting AJs significantly reduced fanning of ARAFs (Fig. 5G, Table S10). Where ARAFs were entirely disconnected from the 1°:LC border, fibers were organized approximately in parallel (Fig. 5B,C, brackets). Hence, connecting the ARAF network to AJs is essential to fan ARAFs appropriately.

Reducing Ecad or α-Cat also generated striking cytoskeletal defects in LCs. Apical F-actin was absent from LCs in which *Ecad^RNAi^* obliterated all AJs. However, in LCs with remnant AJs, large ‘tufts’ of F-actin were observed, usually adjacent to the remaining AJs (Fig. S4A,B). In LCs with *α-cat^RNAi^*, actin radiated from aggregates of cytoplasmic Ecad puncta (Fig. S4C,D). These data suggest that AJs scaffold actin nucleators or regulators in LCs, as has been established in other systems (for example Grikscheit et al., 2015; Kobielak et al., 2004; Kovacs et al., 2002; Kovacs et al., 2011; Magie et al., 2002; Sousa et al., 2005; Tang and Brieher, 2012).

### ARAFs are crosslinked into a network that is connected to the apical plasma membrane

Next, we interrogated the relationship between ARAFs and the apical plasma membrane. Spectrin complexes link F-actin to the cell membrane and, in also cross- linking actin filaments, can introduce order into the cortical actin network (Leterrier and Pullarkat, 2022; Lorenzo et al., 2023). In axons, cortical rings of actin are tethered to the membrane, linked to each other, and periodically spaced by Spectrins. We hypothesized that Spectrins might similarly organize ARAFs and anchor the network to the apical membrane. Spectrins are tetramers, composed of two heterodimers which in *Drosophila* are of α-Spectrin (α-Spec) and β-Spectrin (β-Spec) or βH-Spectrin (encoded by *Karst*, *Kst*) (Byers et al., 1992; Deng et al., 1995; Dubreuil et al., 1987; Dubreuil et al., 1990).

We detected α-Spec and β-Spec at apical and basolateral membranes of all pupal eye cells whilst Kst was enriched at apical membranes (Fig. 6A,B, data not shown). Similar Spectrin distributions have been reported previously in the fly eye and other tissues (Lee et al., 2010; Lee et al., 1997; Thomas and Kiehart, 1994; Thomas and Williams, 1999; Thomas et al., 1998). More Kst was detected in 1° cells than in LCs, and Kst localized along, adjacent to, and between ARAFs and accumulated toward AJs, especially those between 1°s and LCs (Fig. 6C). This suggests that α-Spec/Kst complexes link ARAFs and tether the network to the apical membrane. To test this, we reduced *α-Spec* or *kst* expression (by 29.03% and 60.8% respectively, Table S3), which generated a less dense and randomly organized ARAF network, with small patches devoid of actin (Fig. 6D-G, Table S11). This phenotype is consistent with partial collapse of the ARAF network when not securely connected to the apical membrane and crosslinking is weakened.

**Figure 6:**
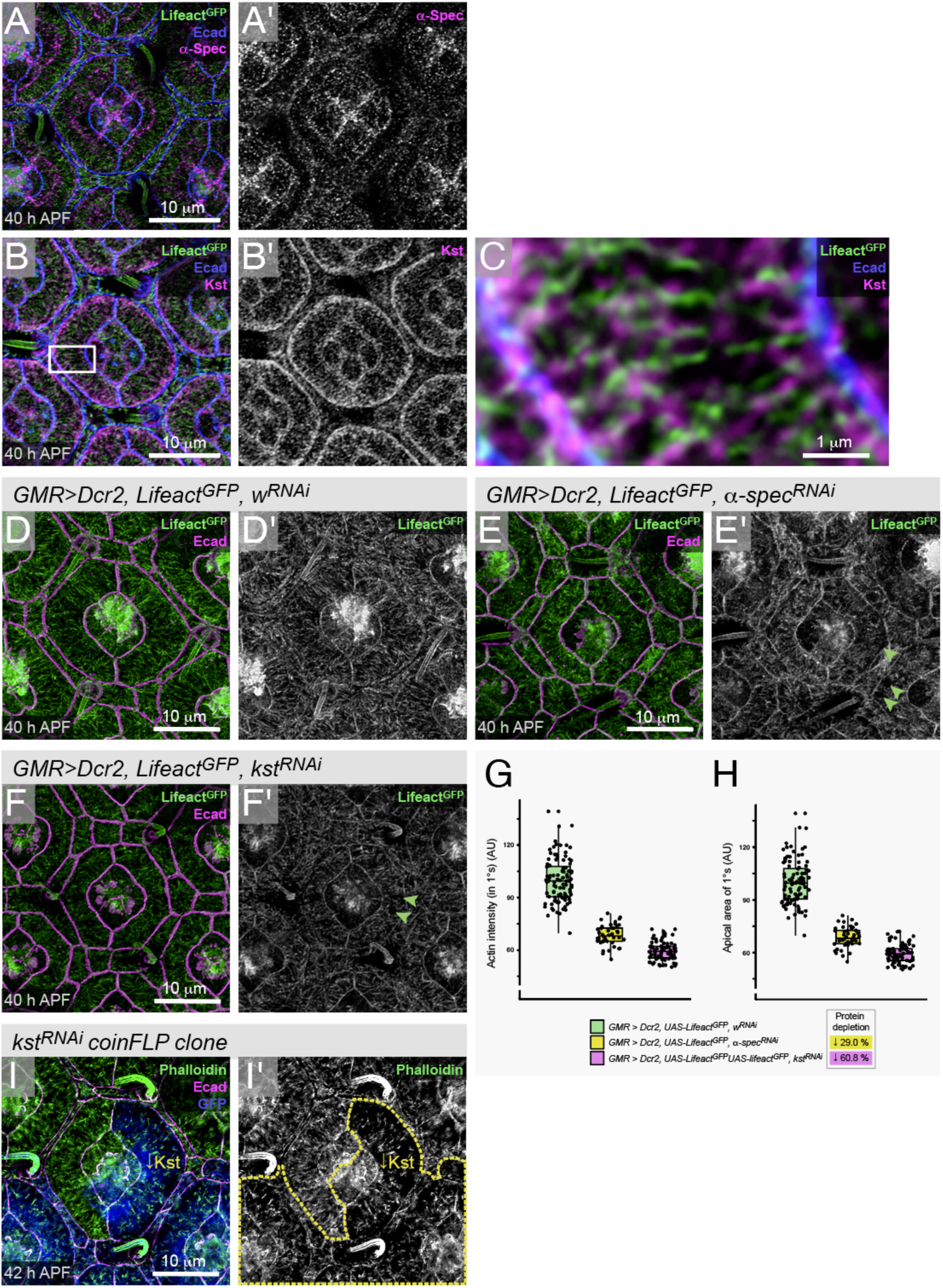
Spectrins connect ARAFs to the cell membrane. (A) α-Spec and (B) Kst, in magenta, detected in ommatidia at 40 h APF. Lifeact labels F-actin and Ecad marks AJs. Boxed region in (B) is shown at higher magnification in (C). (D) A control ommatidium with *w^RNAi^* expression and ommatidia with (E) α-*spec^RNAi^*or (F) *kst^RNAi^* expression. Quantification of (G) the intensity of apical-cortical F-actin and (H) transverse apical area of 1° cells; see Tables S11 and S12 for statistical test scores. (I) A hybrid ommatidium with one 1° cell expressing *kst^RNA^*, labelled with GFP (in blue). F-actin labelled with phalloidin (green). In (I’), yellow dashed line outlines tissue with *kst^RNAi^* expression, marked by GFP. Scale bars are 1μm and 10μm.

We expected that even only partially disconnecting ARAFs from the apical membrane could lead to larger relaxed 1°s, but these cells were instead smaller (Fig. 6E,F,H, Table S12). However, in these experiments LCs were wider which, we speculate resisted expansion of the 1° cell awning (Fig. 6E,F). Indeed, when we used the coinFLP system (Bosch et al., 2015) to generate hybrid ommatidia with just one 1° cell expressing *kst^RNAi^* or α-*Spec^RNAi^*, that 1° cell was consistently larger (Fig. 6I and data not shown). These observations concur with an earlier description of Spectrins in maintaining apical cortical tension to restrict the size of pupal eye cells (Deng et al., 2020). We now show that in 1°s it is specifically the contractile ARAF network that introduces cortical tension that restricts the size of the apical awning.

Villin proteins are primarily actin bundlers, but Villins have also been shown to tether actin to the membrane and sever or nucleate actin, depending on their post-translational modification (Khurana and George, 2008; Sharkova et al., 2023). We found that the *Drosophila* Villin Quail (Qua) (Mahajan-Miklos and Cooley, 1994) was expressed at higher levels in 1°s and accumulated along ARAFs as they formed (Fig. 7B). By 40 h APF, virtually no Qua was observed in LCs, and Qua was highly enriched along ARAFs (Fig. 7C,D). Reducing Qua (by 65.4%, Table S3) rendered the ARAF network severely eroded: remaining actin fibers were short and randomly organized and 1° cells severely misshapen (Fig. 7E-G, Table S13). These data are consistent with collapse of the ARAF network consequent to loss of bundling by Qua, and compromised tethering to the apical membrane. Hence we posit that ARAFs are not individual actin filaments but rather bundles, and we therefore refer to them as fibers.

**Figure 7:**
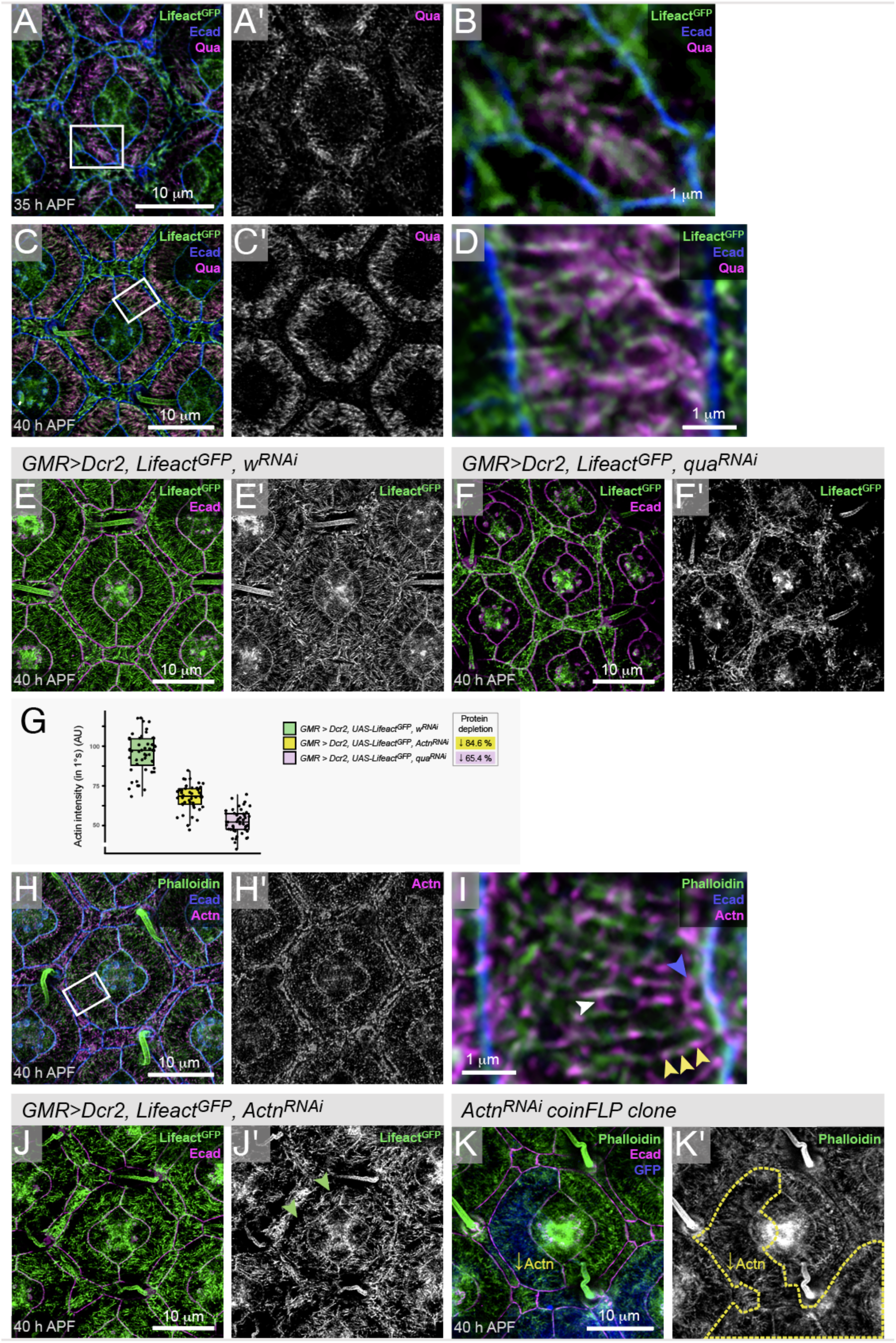
Qua and Actn are crucial for ARAF organization. Detection of Qua (A) at 35 h APF, with boxed area at higher magnification in (B). (C) Qua in an ommatidium at 40 h APF. Boxed area shown at higher magnification in (D). (E) Control *GMR>w^RNAi^* ommatidium, and (F) ommatidia with *qua^RNAi^*expression (G) Analyses of intensity of apical-cortical F-actin density; see Table S13 for statistical test scores. (H) Actn detected in the eye at 40 h APF, with boxed area enlarged in (I). Yellow arrowheads indicate periodic distribution of Actn along an ARAF, white arrowhead indicates Actn detected linking ARAFs, blue arrowhead highlights Actn accumulation toward the AJ. (J) Eye with *Actn^RNAi^* expression, with green arrowheads indicating gaps between ARAF network and the AJ. (K) A hybrid ommatidium with one 1° cell with *Actn^RNAi^*, labelled with GFP (in blue). F-actin labelled with phalloidin (green). Yellow dashed line in (K’), outlines tissue with *Actn^RNAi^*, with GFP (blue). Scale bars are 1μm and 10μm.

Our search for other actin bundling proteins that are expressed in the pupal eye (data not shown) also identified non muscle α-Actinin (Actn) (Fyrberg et al., 1990) which localized to ARAFs at 40 h APF and accumulated on ARAFs toward ZAs (Fig. 7H,I). Our observations are consistent with Actn localization to F-actin, stress fibers, and ZAs in other fly and vertebrate tissues (Lopez-Gay et al., 2020; Pradhan et al., 2023; Rodriguez-Diaz et al., 2008; Wahlstrom et al., 2004), reviewed by Rajan et al., 2023; Sjoblom et al., 2008). Close inspection revealed periodic distribution of Actn along many ARAFs (Fig. 7I, yellow arrowheads) and bridging gaps between neighboring ARAFs (Fig. 7I, white arrowheads). Reducing Actn (by 84.6%) reduced the density of apical cortical actin, and many remaining fibers were shorter, curled, or randomly oriented, consistent with a role for Actn in crosslinking ARAFs to organize and maintain the network (Fig. 7G,J, Table S13). We also observed many small gaps between the remaining ARAF network and the 1° cell boundary (Fig. 7J, green arrowheads), similar to the gaps observed when we reduced Ecad or α-Cat, although far less severe. This suggests that Actn may contribute to linking the ARAF network to ZA-actin or AJs and, consistent with this role, even mildly reducing Actn in one 1° cell of an ommatidium consistently enlarged that 1° (Fig. 7K).

Next, we simultaneously reduced Qua and Actn (Fig. S5). This devastated the ARAF network and severely impeded the rounding of 1°s. These data underscore that cross- linking of actin bundles into an ARAF network, securing the network to the apical membrane, and tethering the network to ZAs, is essential for integrity of the ARAF network and to correctly shape 1° cells.

### ARAFs introduce or support curvature of the apical membrane of 1° cells

Given the three-dimensional nature of the cytoskeleton, we next considered the structure of the ARAF network from this perspective. 3-D renderings of our confocal images revealed that in the *Drosophila* retina, the apical cytoskeleton, and by inference the membrane to which it is tethered, changes dramatically from 35 to 40 h APF (Fig. 8A-E).

**Figure 8:**
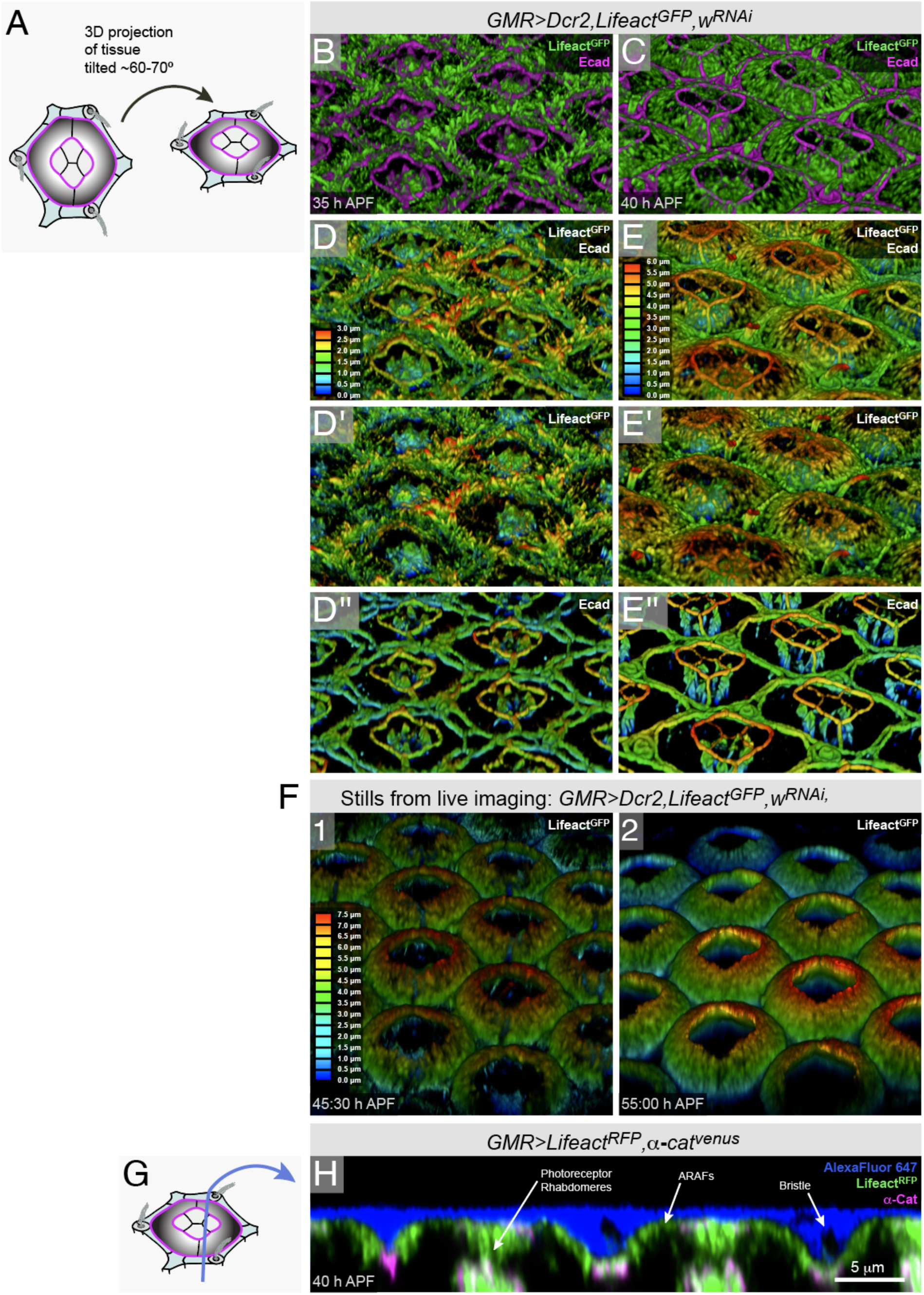
Ommatidia become domed as they mature. (A) Cartoon of ommatidia, with one tilted at roughly the same angle as 3-D perspective images in panels (B) to (F). (B) 3-D perspective images of the eye at 35 h APF and (C) at 40 h APF, with F-actin in green and Ecad in magenta. For (D) and (E), images (B) and (C) are presented with depth color-coding as follows: (D) and (E) both Lifeact^GFP^ and Ecad are shown, (D’) and (E’) only Lifeact^GFP^ shown, (D’) and (E’) only Ecad shown. (F) 3-D depth-coded images of F-actin in the same retina at 45:30 h APF and 55 h APF. Movie S2 documents the ARAF network in these ommatidia from 44 to 55 h APF. (G) Cartoon of ommatidium with blue line indicating plane of longitudinal section of the image presented in (H), a section through the apical region of an eye, with Lifeact^RFP^ and α-Cat detected in the cells and the extracellular mounting medium labelled with AlexaFluor® 647. Scale bar is 5 μm.

Specifically, by 40 h APF, when the ARAF network was well formed, 3-D renderings showed that the fibers arched from the outer ZAs (those between 1° and LCs) to inner ZAs (those between 1° and CCs) (Fig. 8C,E). The arching morphology of fibers is referenced in the name Apical Ribs of Actin Fibers. Our renderings also revealed that numerous filopodial-like structures project apically from 2° and 3° cells at 35 h APF (Fig. 8B,D), but these vanished by 40 h APF (Fig. 8C,E). Hence, retraction of apical filopodia from the LCs, and the formation of ARAFs, transformed the retina into a landscape of repeating ‘ommatidial domes’. After 40 h APF, ARAFs became increasingly dense and domes more pronounced (Fig. 8F, Movie S3). Unfortunately, expression of membrane-tethered tags (e.g. palmitoylated mKate) failed to clearly label the apical membrane of ommatidia, but instead imaging retinas (that had not been exposed to detergent) in mounting medium flooded with AlexaFluor 647 (that remained extracellular) allowed us to confirm that our 3-D renderings of ARAFs reflected the domed shape of the apical surface of ommatidia (Fig. 8G,H). Longitudinal sections of these ommatidia also revealed that the apical membrane of central CCs is relatively flat in comparison to the rounded 1° cells (Fig. 8H).

Concurrent with the formation of ARAFs, we observed that by 40 h APF the inner ZAs of 1° cells were higher than the outer ZAs (Fig. 8C,E). We measured this as elevation (difference between apical-basal placement of inner and outer ZAs of 1°s), which we found increased from 0.55 μm at 35 h APF (SD = 0.13) to 2.29 μm at 40 h APF (SD = 0.31) (Figs 9A, S6, Table S14). To test whether ARAFs were necessary for elevation of these junctions, we impaired the ARAF network by hampering the formation of ARAFs (by reducing Dia, Fig. 9B,D), hampering formation of ZA-actin (by reducing Ena, Fig. 9C,E), or disabling bundling and apical anchorage of ARAFs (by reducing Qua, not shown). In each of these treatments, the inner ZAs remained at the same elevation as that observed in wild type ommatidia at 35 h APF (Figs 9A, S6).

**Figure 9:**
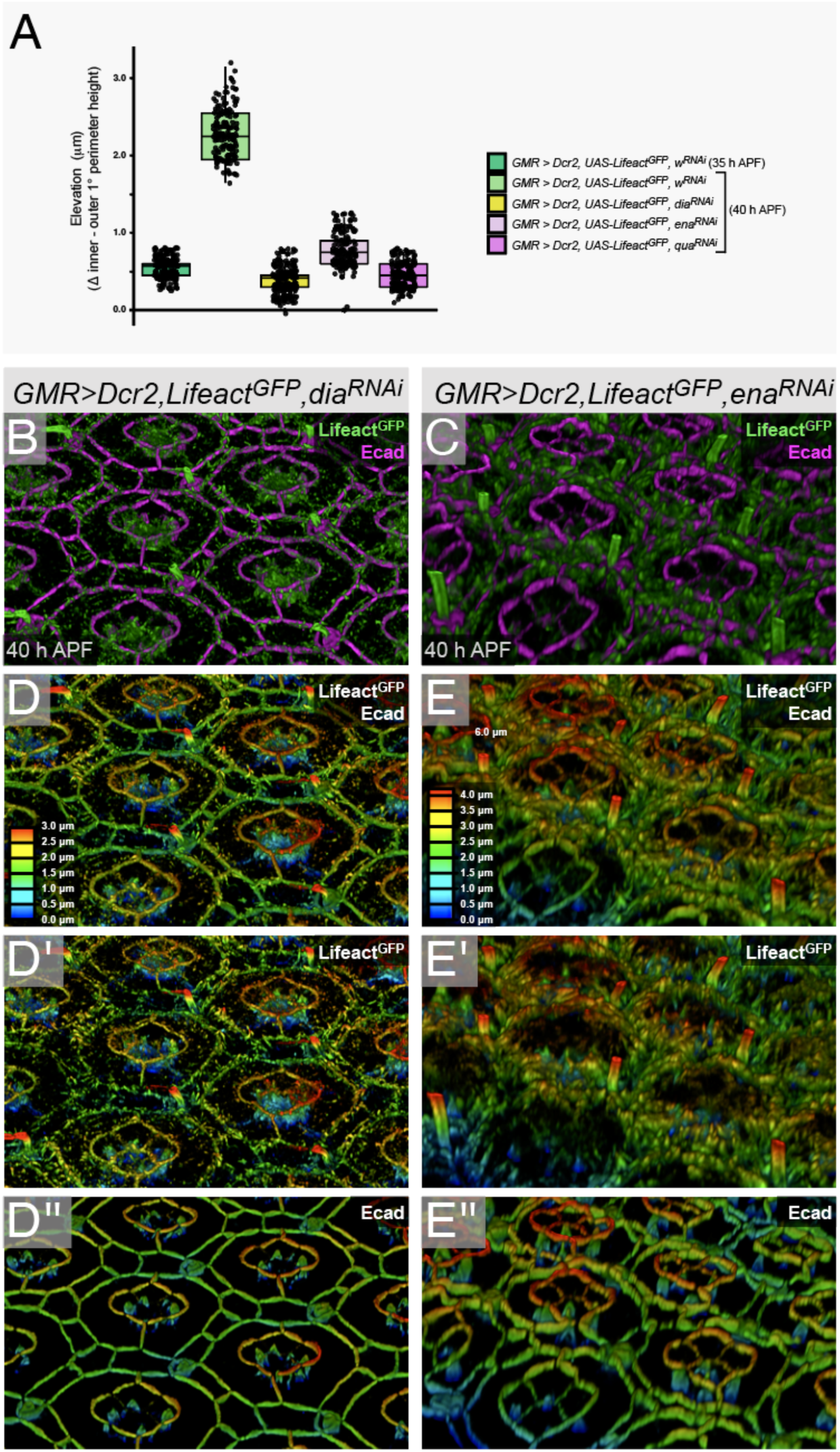
Ommatidial doming is dependent on the ARAF network. (A) Quantification of the elevation of the inner 1°:CC ZAs, in relation to outer 1°:LC ZAs at 35 and 40 h APF, of genotypes as indicated in key. See Fig. S6 for more detailed analyses; and Table S13 for statistical test scores. (B) 3-D perspective image of an eye expressing *dia^RNAi^* or (C) *ena^RNAi^*at 40 h APF. F-actin in green and Ecad in magenta. For (D) and (E), images (B) and (C) are presented with depth color-coding as follows: (D) and (E) both Lifeact^GFP^ and Ecad are shown, (D’) and (E’) only Lifeact^GFP^ shown, (D’) and (E’) only Ecad shown. Refer to Fig. 8 (C) and (E) for control eye with *w^RNAi^*expression.

Taken together, we conclude that the ARAF network is built to drive upward expansion and doming of ommatidia. During this morphogenesis, the inner 1° cell junctions move apically, movement that is either driven or supported by the ARAFs. Earlier, we described that ARAFs constrain the transverse expansion of the 1° cell awning and ensure that 1°s attain their characteristic fabiform shapes. Hence, the ARAFs exert morphogenetic forces in both transverse and longitudinal axes.

Finally, to better understand how the ARAF network shapes 1° cells, we built 3-D renderings of 1° cells from z-stacks that captured α-Spec at all membranes at 40 h APF (Fig. 10A-C, Movies S3, S4 and S5). These renderings revealed that 1° cells have a complex shape not captured in traditional 2-D drawings of ommatidia (Fig. 1A). The stereotypical fabiform shape is generated by widening only the apical region of 1° cells into an awning, which reaches over and anchors to CCs. Tensile ARAFs span the width of this awning (Fig. 10B,D). The top third of each 1° connects to a partner 1° to form a collar around the CCs. The basal third of each 1° cell narrows and projects toward the base of the ommatidium.

**Figure 10:**
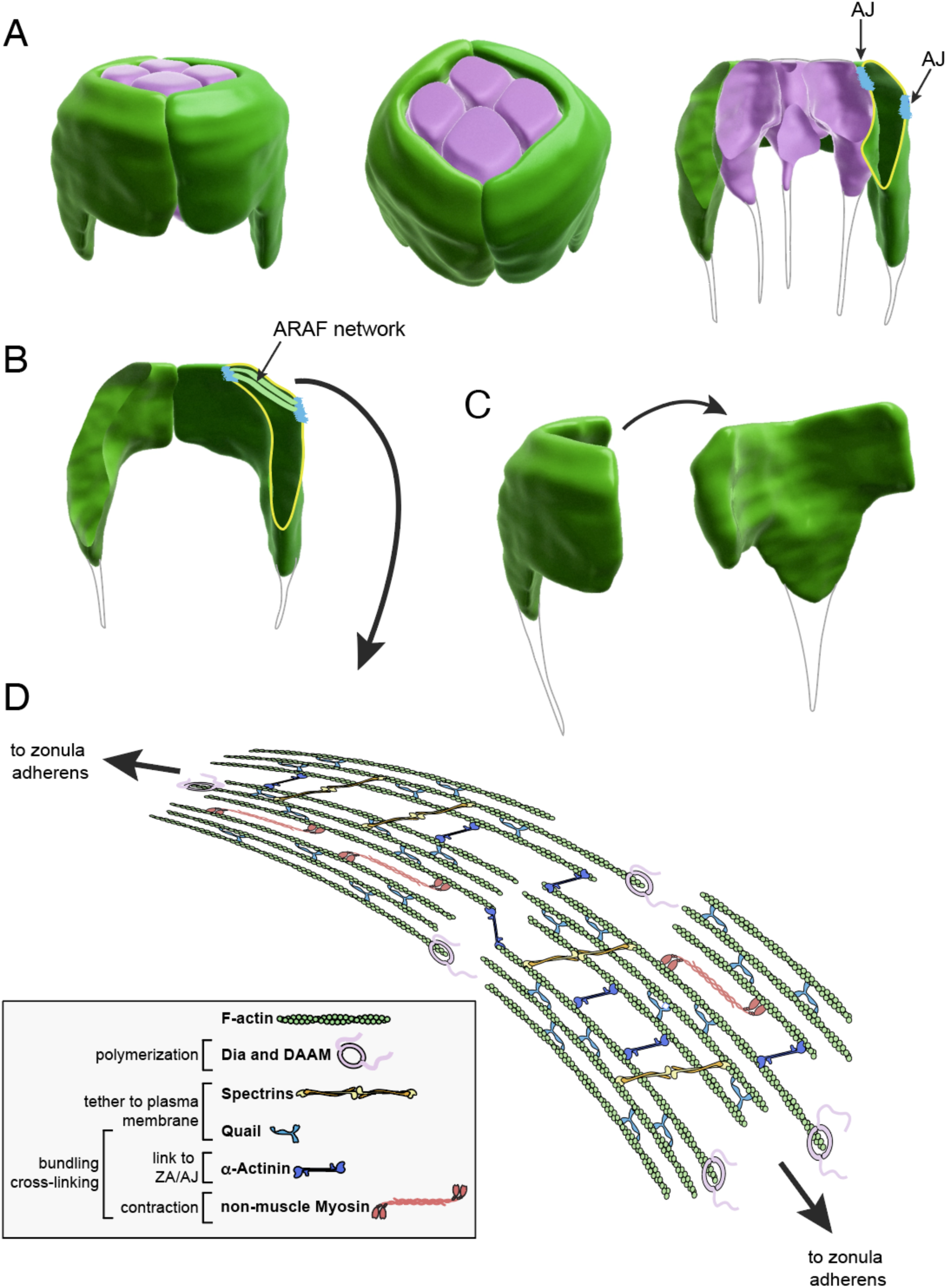
ARAFs support the complex shape of 1° cells. (A) 3-D rendering of the cone and 1° cells of an ommatidium. For the image at right, the ommatidium was bisected along the midline; cleavage of the left 1° is indicated in yellow and AJs drawn in blue. (B) A pair of 1°s, with ARAFs and AJs annotated. (D) A single 1° at two orientations. Basal projections of the cells are annotated since rendering these was not successful. See movies S3 to S5. (D) Model of the ARAF network, with proteins responsible for its construction and maintenance.

## Discussion

We have found that the apical region of 1° cells becomes filled with actin fibers organized into a complex ARAF network (Figs 1, 2, and 8). Meanwhile, in 3° and horizontal 2° cells a ‘starburst’ organization of actin was observed, and no obvious actin organization was discernible in oblique 2° cells. Hence, cell-type and shape-specific actin structures emerge in the *Drosophila* pupal eye as the tissue matures. Each of these cells occupy distinct positions within the eye, and we predict that the cells’ cytoskeletal architecture is in large part the mechanical response to position-dependent forces transmitted from cell neighbors. However, our interrogation of 1° cells indicates that the mechanical response and/or the cytoskeletal structure that is built in response, can be pre-determined by genetic control.

Higher resolution imaging is required to confirm the precise structure of the ARAF network. However, our data supports a model where ARAFs are composed of actin bundles that connect in tandem to span the width of 1°s, and where ARAFs connect laterally, to generate an organized network (Fig. 10D). Our 3-D renderings of the network (Figs 8 and 9), show that ARAFs are curved and that the network arches upward from the outer ZAs (between 1°s and LCs) toward the inner ZAs (between 1° and CCs). ARAFs occupy only the apical region of 1° cells, a region that is enlarged and bends to generate an ‘awning’ that partially covers the neighboring CCs. Compromising the formation or maintenance of ARAFs, or their connection to 1° cell ZAs, led to flattened 1°s with awnings that failed to acquire their consistent width and rounded shape (Figs 4-7, 9). Hence the ARAF network is essential for ommatidia to be correctly shaped.

We identified a comprehensive set of proteins that assemble and maintain the ARAF network (Fig. 10D). The formins Dia and DAAM function together to assemble the filaments of the ARAF network. This conclusion is supported by the presence of Dia and DAAM at the tips of ARAFs as the network emerged (35 h APF, Fig. 3), and severe loss of ARAFs when Dia and/or DAAM activity were compromised (Fig. 4). It is not clear how, precisely, the ARAFs are initiated, and it is plausible that elements of the apical-medial actin meshwork present in younger 1° cells (prior to ∼31 h APF) are reorganized to form the first ARAFs or severed to generate short polymers from which ARAFs are nucleated. Alternatively, as has been described for actin cable formation in nurse cells (Logan et al., 2022), the fibers are generated entirely *de novo* by membrane-anchored formins.

Dia, and later DAAM, were enriched at AJs and these could be principal sites from which new ARAFs form, but Dia and DAAM localized through the apical cortex where they associated with actin tips (Fig. 3). Hence, our model assumes that ARAFs are nucleated throughout the apical cortex of 1°s and that bundling and crosslinking reorient the fibers into an organized network that spans the 1° cell width.

ARAFs are linked to the apical membrane and to each other (Fig. 10D). This statement is supported by our detection of Spectrins and the villin Qua along nascent and maturing ARAFs and detection of Actn along and between ARAFs (Figs 6 and 7). Spectrins function primarily to tether actin to plasma membranes, but they also link actin filaments to each other (Leterrier and Pullarkat, 2022; Lorenzo et al., 2023). Vilins are primarily actin bundlers, but they can also link actin to membranes (Khurana and George, 2008; Sharkova et al., 2023). Actns are actin crosslinkers that also link F-actin to junctions (Rajan et al., 2023; Sjoblom et al., 2008). Our model hence posits that these proteins function in these ways to either assemble or maintain the ARAF network, but more detailed analyses are needed to test these assumptions. Nonetheless, reducing the expression of each of these proteins generated phenotypes consistent with a failure to bundle F-actin into ARAFs (*qua^RNAi^* expression), a failure to link neighboring or tandem ARAFs to each other (*Actn^RNAi^*), a failure to link ARAFs to AJs (*Actn^RNAi^*) and the disconnection of ARAFs from the apical membrane (*qua^RNAi^*, *kst^RNAi^*or *α-spec^RNAi^*) (Figs 6 and 7). The severity of each of the phenotypes we observed would have been dependent on the degree to which the target proteins were reduced and the extent to which the ARAF network could be maintained by non-targeted proteins.

Our rendering of the 3-D shape of the ARAF network crucially revealed that the network is curved and arches from the outer AJs between 1° and LCs, upward toward inner 1°:CCs AJs (Figs 8, 9A). Using localization of α-Spec as a proxy for the cell membrane, we built 3-D renderings of 1° cells that confirmed that the apical membranes are curved to generate domed ommatidia (Fig. 10). Hence, the ARAFs are like rafters of a roof: periodically spaced and supporting and shaping the membrane above. Curiously, disrupting ARAFs led to a change in the placement of the inner AJs: these were considerably lower than in wild type ommatidia, and the ommatidia were consequently ‘pancaked’ not only due to disruption of the rafter-like ARAFs, but also because the inner AJs that connect 1°s to CCs were not elevated.

The ARAF network is linked to the ZAs of 1° cells (Fig. 10D). This statement stems from observations that ARAFs synapse with ZA-actin or AJs (Figs 1 and 2), and multiple genetic manipulations support this interpretation. First, ablating AJs by reducing Ecad or α-Cat ‘unhooked’ the ARAF network from the 1° cell membrane, but ARAFs remained connected to the cell border where AJs remained (Fig. 5). Second, reducing Actn introduced gaps between the ARAF network and ZAs (Fig. 7), consistent with Actn participation in linking actin to AJs as established in other systems (Knudsen et al., 1995; Nieset et al., 1997). Third, reducing Ena, which is required for actin assembly at ZAs (Baum and Perrimon, 2001; Leerberg et al., 2014; Scott et al., 2006), introduced disorganization to the ARAF network (Figs 3 and 4), underscoring that the ARAF network must be hooked to a stable ZA to maintain its organization.

Decoupling the ARAF network from AJs provided strong evidence that ARAFs are tensile apical stress fibers. Specifically, our reduction of Ecad or α-Cat left AJs connecting 1°:CCs intact with ARAFs attached, but the ARAFs disconnected from the outer AJs and retracted into a rounded shape that mimicked the fabiform shape of the 1° cell awning (Fig. 5B,C). Further, the disconnected outer membranes of 1°s billowed outward (Fig. 5B,C). These experiments, which are a genetic equivalent to laser ablation, convey that a critical role for the tensile ARAF network is to limit the girth of 1°s and shape each pair of 1°s into an almost-perfect ring. Returning to our analogy of ARAFs in supporting a house’s roof, the ARAFs are hence like rafter ties that prevent the walls of a house from splaying outward.

Apical stress fibers have been described in two other *Drosophila* tissues – the thoracic and tracheal epithelia. Apical stress fibers in the thorax form in response to pulling mechanical forces (Lopez-Gay et al., 2020). We propose that mechanical forces similarly induce ARAF formation, and changes in the ommatidial core (e.g. elongation of photoreceptors) or narrowing of the LC lattice could provide the mechanical force trigger. Recent modeling suggested that in tracheal epithelial cells the stress fiber network is self-assembled from small ‘clusters’ of F-actin (nanoclusters) in response to anisotropic cortical stress (Sekine et al., 2024). In this model, the availability of actin nanoclusters and crosslinkers, and the rate of actin turnover, governed the morphology of the stress fiber network. Similar rules likely govern how the ARAF network is shaped and our illustration of the network in Figure 10D attempts to convey some of the geometry that we expect fine-scale analyses will reveal. According to a self-assembly model, the organization of ARAFs is dependent on protein availability and activity.

Indeed, whilst we observed Dia, DAAM, activated Rho, Ena, and Actn in 1°s and LCs (Figs 3, 7H, S3), we observed higher expression of Kst and Qua in 1°s (Figs 6B, 7A-D). This suggests that 1° cells are ‘primed’ for the assembly of the ARAF network. How Kst and Qua are regulated to become enriched in 1°s has not been explored.

Our study underscores the importance of understanding the 3-D morphology of the cytoskeleton and cells. The shape of 1° cells has for many years been considered only from the transverse perspective of the apical region. However, our work shows that these are not simple fabiform cells that pair to form neat rings around CCs. Instead, the top third of each 1° cell is expanded and the bottom two-thirds narrows to project to the base of the ommatidium. The ARAFs fill the apical third of 1°s to support and maintain their uniform fabiform shape and the elevated apical membrane that generates domed ommatidia. We predict that similarly extraordinary actin conformations will become evident in other tissues as the 3-D morphologies of other cells are explored.

## Materials and Methods

### Drosophila stocks and genetics

All crosses and *Drosophila* lines were maintained at 25°C. *Drosophila* lines used are listed in Tables S15 and S16. F-actin was detected in retinas of *w^1118^*, or *ubi-lifeact^YFP^* (Santa-Cruz Mateos et al., 2020), or retinas with *UAS-Lifeact^GFP^*, *UAS-Lifeact^RFP^*or *UAS-GMA* (Bloor and Kiehart, 2001; Hatan et al., 2011; Huelsmann et al., 2013) driven by *GMR-GAL4* which initiates expression in eye cells after passage of the morphogenetic furrow (Freeman, 1996). *GMR-GAL4, UAS-Lifeact^GFP^*, or *GMR-GAL4, UAS-Lifeact^RFP^* or *UAS-Dcr2; GMR-GAL4, UAS-Lifeact^GFP^* (abbreviated to *GMR>Lifeact^GFP^*, and *GMR>Lifeact^RFP^*, and *UAS-Dcr2; GMR> Lifeact^GFP^*respectively) were crossed to UAS transgenics to reduce proteins of interest (see below). The coinFLP system was used to generate clones of eye cells with *UAS-RNAi* transgene expression (Bosch et al., 2015).

### RNA interference

To target genes of interest, all *UAS-RNAi* transgenes available from the Bloomington *Drosophila* Stock Center and Vienna *Drosophila* Resource Center at commencement of this project were obtained (Table S16) with the exception of RNAi transgenes for *Ecad* and *α-cat* (Seppa et al., 2008). Transgenics were crossed to *GMR>Lifeact^GFP^* and *UAS- Dcr2; GMR> Lifeact^GFP^*, and pupal eyes of progeny assayed for disruption to the ARAF network and associated eye mis-patterning. For RNAi transgenes targeting the same loci, all pupal eye phenotypes were qualitatively similar, and transgenes that most severely disrupted ARAFs were selected for experiments presented in this work. Degree of protein knockdown by RNAi transgenes was determined by detecting the target protein with immunofluorescence in retinas with the associated transgene, and comparing to control tissue expressing a transgene against *white* (*w*) (Table S3). The actin cytoskeleton was unperturbed by *w^RNAi^* or *Dcr2* expression. Occasional errors in the placement or number of LCs are observed in retinas heterozygous for *GMR-GAL4*.

### Eye dissection and immunohistochemistry

Pre-pupae were gathered and incubated at 25°C in humidity chambers until dissection as previously described (DeAngelis and Johnson, 2019) but with the following adaptations to preserve the cytoskeleton: First, after dissection in ice-cold phosphate buffered saline (PBS), eye-brain complexes were fixed in 4% formaldehyde in PBS supplemented with phalloidin (1:500) on ice for 35 minutes. Second, tissue was maintained on ice or at 4°C whenever possible. Third, to preserve the shape of ommatidia and the ARAF network, eyes were mounted on slides surrounded by a human hair that raised the coverslip from the retinas. Fourth, retinas were imaged within a few hours of preparation.

For ‘rapid imaging’, after fixation eyes, were rinsed in ice-cold PBS and PBT (PBS with 0.15% Triton-X), incubated in mounting medium (0.5% n-propyl gallate in 80% glycerol in PBS) for 1 hour, mounted on slides, and imaged immediately. Alternatively, after fixation and rinsing, eyes were dissected from optic lobes on slides, excess PBT removed, and mounting medium applied directly to the tissue, which was cover-slipped and immediately imaged.

For immunofluorescence, after fixation and rinsing, retinas were incubated with antibodies listed in Table S17. Endogenous fluorescence was detected when imaging Actn^GFP^, DAAM^GFP^, GMA, Kst^GFP^, Lifeact^GFP^, Lifeact^RFP^, Lifeact^YFP^, shg^mTom^ and α-Spec^GFP^ (Table S15). When used for F-actin detection, rhodamine-phalloidin was included in the primary antibody solution (1:500, Thermo Fisher Scientific, cat# R415).

Live imaging was performed as previously described (DeAngelis and Johnson, 2019), with the following modifications: pupae were positioned with their abdomens and head on a support of soft paper putty, and surrounded by a circle of putty onto which the cover slip was placed. The cover slip was depressed to reach but not touch the eye.

### Protein localization

Localization/expression of ARAF-associated proteins was assayed with primary antibodies (Table S17) and confirmed by GFP-fusion transgenics or reporter transgenes if available (Table S15). Immunofluorescence was compared in retinas with and without RNAi transgenes targeting the detected protein (Table S16, data not shown).

### Image acquisition and processing

High -esolution Z-stack images were acquired with a Leica SP8 confocal microscope and proprietary Leica Application Suite X software (LASX) and LIGHTNING deconvolution. Images were gathered with a scan format of 2048 x 2048 and resolution of 45 nm x 45 nm per pixel. Control and experimental retinas were imaged using precisely the same parameters. Maximum projections of sections of the apical region of 1° cells were generated in LASX and processed in Adobe Photoshop®, with control and experimental images processed identically. To visualize ARAF architecture, Z-stacks of Lifeact^GFP^ were rendered as 3-D and depth-coded images using the 3-D feature of LASX. To determine the 3-D architecture of 1° cells, Z-stacks of α-Spec immunofluorescence were rendered as 3-D images using the surface feature of Imaris (Oxford Instruments).

### Image analysis and statistics

The polygon tool of Fiji (Schindelin et al., 2012) was used to trace the apical outlines of 1° cells in which Ecad, Dlg, or ZA-actin were detected, and the area of these regions then measured. For control retinas, measurements were normalized to generate mean values of 100 arbitrary units (AU). For experimental data, each 1° cell area measurement was normalized against the mean area of control 1°s.

After outlining 1°s with Fiji’s polygon tool, actin density was determined as mean intensity of Lifeact^GFP^. To measure ZA-actin density, Fiji’s polygon tool was used to trace the outline of 1°:LC AJs that had been detected with Ecad immunofluorescence. Mean intensity of Lifeact^GFP^ within these outlines was then determined. For controls, mean intensity measurements were normalized to generate a mean of 1 AU and experimental measurements were normalized against the mean actin intensities of controls.

In genotypes where ARAFs were clearly evident, ARAF density was measured by drawing six 2 μm lines per 1°, parallel to the outer perimeter of 1° cells. Fiji’s plot profile function was then used to generate plots of fluorescence intensity of Lifeact^GFP^, Lifeact^RFP^, Lifeact^YFP^ or rhodamine-phalloidin, along each line, with each plot peak representing an intersected actin filament. ARAFs/µm was then calculated.

To determine the orientation of ARAFs with respect to the 1°:LC boundary, Fiji’s angle tool was used to trace individual filaments and the adjacent outer 1° cell ZA, and angles then measured. Neighboring filaments were similarly traced and the angle tool used to measure the angle of orientation between filaments within the ARAF network.

To determine how severely RNAi transgenes reduced target proteins (Table S3), fluorescence intensity of the protein, detected with immunofluorescence, was determined in images of retinas with and without transgene expression, in Fiji. Protein levels were evaluated only in 1° cells.

Elevation of ommatidial domes was calculated as the difference in the position of the confocal z-section that captured the top of AJs between 1° and CCs, and position of the z-section that captured the 1° to LC AJs. Pooled elevation data are presented in Fig. 8B, detailed analyses in Fig. S6.

For all analyses, statistically significant differences between genotypes were determined using the student’s t-tests; p-values and sample sizes are listed in supplementary tables. All graphs were generated using R software.

## Acknowledgments

We thank Maria Martin-Bermudo, Jozsef Mihaly, Claire Thomas and Steven Wasserman for generous gifts of *Drosophila* lines or antibodies; the Bloomington Drosophila Stock Center (www.bdsc.indiana.edu) (NIH P40OD018537) and Vienna Drosophila Resource Center (www.vdrc.at) for numerous stocks; and the Developmental Studies Hybridoma Bank (www.dshb.biology.uiowa.edu) for monoclonal antibodies. This work was supported by NIH 2R15GM114729. Wesleyan University provided additional support for undergraduate summer research.

## Notes

### Competing Interest Statement

The authors have declared no competing interest.

## References

1. Abreu-Blanco, M.T., Verboon, J.M. and Parkhurst, S.M., 2014. Coordination of Rho family GTPase activities to orchestrate cytoskeleton responses during cell wound repair. Curr Biol. 24, 144–155.

2. Afshar, K., Stuart, B. and Wasserman, S.A., 2000. Functional analysis of the Drosophila diaphanous FH protein in early embryonic development. Development. 127, 1887–97.

3. Anhezini, L., Saita, A.P., Costa, M.S., Ramos, R.G. and Simon, C.R., 2012. Fhos encodes a Drosophila Formin-like protein participating in autophagic programmed cell death. Genesis. 50, 672–84.

4. Bai, S.W., Herrera-Abreu, M.T., Rohn, J.L., Racine, V., Tajadura, V., Suryavanshi, N., Bechtel, S., Wiemann, S., Baum, B. and Ridley, A.J., 2011. Identification and characterization of a set of conserved and new regulators of cytoskeletal organization, cell morphology and migration. BMC Biol. 9, 54.

5. Baum, B. and Perrimon, N., 2001. Spatial control of the actin cytoskeleton in Drosophila epithelial cells. Nat Cell Biol. 3, 883–90.

6. Blackie, L., Walther, R.F., Staddon, M.F., Banerjee, S. and Pichaud, F., 2020. Cell-type-specific mechanical response and myosin dynamics during retinal lens development in Drosophila. Mol Biol Cell. 31, 1355–1369.

7. Blanchard, G.B., Etienne, J. and Gorfinkiel, N., 2018. From pulsatile apicomedial contractility to effective epithelial mechanics. Curr Opin Genet Dev. 51, 78–87.

8. Blanchard, G.B., Murugesu, S., Adams, R.J., Martinez-Arias, A. and Gorfinkiel, N., 2010. Cytoskeletal dynamics and supracellular organisation of cell shape fluctuations during dorsal closure. Development. 137, 2743–52.

9. Bloor, J.W. and Kiehart, D.P., 2001. zipper Nonmuscle myosin-II functions downstream of PS2 integrin in Drosophila myogenesis and is necessary for myofibril formation. Dev Biol. 239, 215–28.

10. Booth, A.J.R., Blanchard, G.B., Adams, R.J. and Roper, K., 2014. A dynamic microtubule cytoskeleton directs medial actomyosin function during tube formation. Dev Cell. 29, 562–576.

11. Bosch, J.A., Tran, N.H. and Hariharan, I.K., 2015. CoinFLP: a system for efficient mosaic screening and for visualizing clonal boundaries in Drosophila. Development. 142, 597–606.

12. Breitsprecher, D. and Goode, B.L., 2013. Formins at a glance. J Cell Sci. 126, 1–7.

13. Burkel, B.M., von Dassow, G. and Bement, W.M., 2007. Versatile fluorescent probes for actin filaments based on the actin-binding domain of utrophin. Cell Motil Cytoskeleton. 64, 822–32.

14. Byers, T.J., Brandin, E., Lue, R.A., Winograd, E. and Branton, D., 1992. The complete sequence of Drosophila beta-spectrin reveals supra-motifs comprising eight 106- residue segments. Proc Natl Acad Sci U S A. 89, 6187–91.

15. Cagan, R.L. and Ready, D.F., 1989. The emergence of order in the Drosophila pupal retina. Dev Biol. 136, 346–62.

16. Cai, D., Chen, S.C., Prasad, M., He, L., Wang, X., Choesmel-Cadamuro, V., Sawyer, J.K., Danuser, G. and Montell, D.J., 2014. Mechanical feedback through E-cadherin promotes direction sensing during collective cell migration. Cell. 157, 1146–59.

17. Carramusa, L., Ballestrem, C., Zilberman, Y. and Bershadsky, A.D., 2007. Mammalian diaphanous-related formin Dia1 controls the organization of E-cadherin-mediated cell-cell junctions. J Cell Sci. 120, 3870–82.

18. Casas-Tinto, S. and Portela, M., 2019. Cytonemes, Their Formation, Regulation, and Roles in Signaling and Communication in Tumorigenesis. Int J Mol Sci. 20.

19. Castrillon, D.H. and Wasserman, S.A., 1994. Diaphanous is required for cytokinesis in Drosophila and shares domains of similarity with the products of the limb deformity gene. Development. 120, 3367–77.

20. Cetera, M., Ramirez-San Juan, G.R., Oakes, P.W., Lewellyn, L., Fairchild, M.J., Tanentzapf, G., Gardel, M.L. and Horne-Badovinac, S., 2014. Epithelial rotation promotes the global alignment of contractile actin bundles during Drosophila egg chamber elongation. Nat Commun. 5, 5511.

21. Charlton-Perkins, M.A., Friedrich, M. and Cook, T.A., 2021. Semper’s cells in the insect compound eye: Insights into ocular form and function. Dev Biol. 479, 126–138.

22. Coravos, J.S., Mason, F.M. and Martin, A.C., 2017. Actomyosin Pulsing in Tissue Integrity Maintenance during Morphogenesis. Trends Cell Biol. 27, 276–283.

23. Courtemanche, N., 2018. Mechanisms of formin-mediated actin assembly and dynamics. Biophys Rev. 10, 1553–1569.

24. David, D.J., Tishkina, A. and Harris, T.J., 2010. The PAR complex regulates pulsed actomyosin contractions during amnioserosa apical constriction in Drosophila. Development. 137, 1645–55.

25. Davis, J.D. and Wypych, T.P., 2021. Cellular and functional heterogeneity of the airway epithelium. Mucosal Immunol. 14, 978–990.

26. DeAngelis, M.W. and Johnson, R.I., 2019. Dissection of the Drosophila Pupal Retina for Immunohistochemistry, Western Analysis, and RNA Isolation. J Vis Exp.

27. Delon, I. and Brown, N.H., 2009. The integrin adhesion complex changes its composition and function during morphogenesis of an epithelium. J Cell Sci. 122, 4363–74.

28. Deng, H., Lee, J.K., Goldstein, L.S. and Branton, D., 1995. Drosophila development requires spectrin network formation. J Cell Biol. 128, 71–9.

29. Deng, H., Yang, L., Wen, P., Lei, H., Blount, P. and Pan, D., 2020. Spectrin couples cell shape, cortical tension, and Hippo signaling in retinal epithelial morphogenesis. J Cell Biol. 219.

30. Dubreuil, R., Byers, T.J., Branton, D., Goldstein, L.S. and Kiehart, D.P., 1987. Drosophilia spectrin. I. Characterization of the purified protein. J Cell Biol. 105, 2095–102.

31. Dubreuil, R.R., Byers, T.J., Stewart, C.T. and Kiehart, D.P., 1990. A beta-spectrin isoform from Drosophila (beta H) is similar in size to vertebrate dystrophin. J Cell Biol. 111, 1849–58.

32. Dutta, D., Bloor, J.W., Ruiz-Gomez, M., VijayRaghavan, K. and Kiehart, D.P., 2002. Real-time imaging of morphogenetic movements in Drosophila using Gal4-UAS-driven expression of GFP fused to the actin-binding domain of moesin. Genesis. 34, 146–51.

33. Edwards, K.A., Demsky, M., Montague, R.A., Weymouth, N. and Kiehart, D.P., 1997. GFP-moesin illuminates actin cytoskeleton dynamics in living tissue and demonstrates cell shape changes during morphogenesis in Drosophila. Dev Biol. 191, 103–17.

34. Eenjes, E., Tibboel, D., Wijnen, R.M.H. and Rottier, R.J., 2022. Lung epithelium development and airway regeneration. Front Cell Dev Biol. 10, 1022457.

35. Emmons, S., Phan, H., Calley, J., Chen, W., James, B. and Manseau, L., 1995. Cappuccino, a Drosophila maternal effect gene required for polarity of the egg and embryo, is related to the vertebrate limb deformity locus. Genes Dev. 9, 2482–94.

36. Faix, J. and Rottner, K., 2022. Ena/VASP proteins in cell edge protrusion, migration and adhesion. J Cell Sci. 135.

37. Fernandez-Gonzalez, R. and Harris, T.J.C., 2023. Contractile and expansive actin networks in Drosophila: Developmental cell biology controlled by network polarization and higher-order interactions. Curr Top Dev Biol. 154, 99–129.

38. Fernandez-Gonzalez, R. and Zallen, J.A., 2011. Oscillatory behaviors and hierarchical assembly of contractile structures in intercalating cells. Phys Biol. 8, 045005.

39. Freeman, M., 1996. Reiterative use of the EGF receptor triggers differentiation of all cell types in the Drosophila eye. Cell. 87, 651–60.

40. Fyrberg, E., Kelly, M., Ball, E., Fyrberg, C. and Reedy, M.C., 1990. Molecular genetics of Drosophila alpha-actinin: mutant alleles disrupt Z disc integrity and muscle insertions. J Cell Biol. 110, 1999–2011.

41. Gertler, F.B., Comer, A.R., Juang, J.L., Ahern, S.M., Clark, M.J., Liebl, E.C. and Hoffmann, F.M., 1995. enabled, a dosage-sensitive suppressor of mutations in the Drosophila Abl tyrosine kinase, encodes an Abl substrate with SH3 domain-binding properties. Genes and Development. 9, 521–33.

42. Grikscheit, K., Frank, T., Wang, Y. and Grosse, R., 2015. Junctional actin assembly is mediated by Formin-like 2 downstream of Rac1. J Cell Biol. 209, 367–76.

43. Grosshans, J., Wenzl, C., Herz, H.M., Bartoszewski, S., Schnorrer, F., Vogt, N., Schwarz, H. and Muller, H.A., 2005. RhoGEF2 and the formin Dia control the formation of the furrow canal by directed actin assembly during Drosophila cellularisation. Development. 132, 1009–20.

44. Hannezo, E., Dong, B., Recho, P., Joanny, J.F. and Hayashi, S., 2015. Cortical instability drives periodic supracellular actin pattern formation in epithelial tubes. Proc Natl Acad Sci U S A. 112, 8620–5.

45. Harris, T.J.C., 2018. Sculpting epithelia with planar polarized actomyosin networks: Principles from Drosophila. Semin Cell Dev Biol. 81, 54–61.

46. Hatan, M., Shinder, V., Israeli, D., Schnorrer, F. and Volk, T., 2011. The Drosophila blood brain barrier is maintained by GPCR-dependent dynamic actin structures. J Cell Biol. 192, 307–19.

47. He, L., Wang, X., Tang, H.L. and Montell, D.J., 2010. Tissue elongation requires oscillating contractions of a basal actomyosin network. Nat Cell Biol. 12, 1133–42.

48. Hewitt, R.J. and Lloyd, C.M., 2021. Regulation of immune responses by the airway epithelial cell landscape. Nat Rev Immunol. 21, 347–362.

49. Homem, C.C. and Peifer, M., 2008. Diaphanous regulates myosin and adherens junctions to control cell contractility and protrusive behavior during morphogenesis. Development. 135, 1005–18.

50. Huelsmann, S., Ylanne, J. and Brown, N.H., 2013. Filopodia-like actin cables position nuclei in association with perinuclear actin in Drosophila nurse cells. Dev Cell. 26, 604–15.

51. Jacinto, A., Wood, W., Balayo, T., Turmaine, M., Martinez-Arias, A. and Martin, P., 2000. Dynamic actin-based epithelial adhesion and cell matching during Drosophila dorsal closure. Curr Biol. 10, 1420–6.

52. Jiao, R., Daube, M., Duan, H., Zou, Y., Frei, E. and Noll, M., 2001. Headless flies generated by developmental pathway interference. Development. 128, 3307–19.

53. Johnson, H.W. and Schell, M.J., 2009. Neuronal IP3 3-kinase is an F-actin-bundling protein: role in dendritic targeting and regulation of spine morphology. Mol Biol Cell. 20, 5166–80.

54. Johnson, R.I., 2021. Hexagonal patterning of the Drosophila eye. Dev Biol. 478, 173–182.

55. Khurana, S. and George, S.P., 2008. Regulation of cell structure and function by actin-binding proteins: villin’s perspective. FEBS Lett. 582, 2128–39.

56. Kiehart, D.P., Galbraith, C.G., Edwards, K.A., Rickoll, W.L. and Montague, R.A., 2000. Multiple forces contribute to cell sheet morphogenesis for dorsal closure in Drosophila. J Cell Biol. 149, 471–90.

57. Knudsen, K.A., Soler, A.P., Johnson, K.R. and Wheelock, M.J., 1995. Interaction of alpha-actinin with the cadherin/catenin cell-cell adhesion complex via alpha-catenin. J Cell Biol. 130, 67–77.

58. Kobielak, A., Pasolli, H.A. and Fuchs, E., 2004. Mammalian formin-1 participates in adherens junctions and polymerization of linear actin cables. Nat Cell Biol. 6, 21–30.

59. Kovacs, E.M., Goodwin, M., Ali, R.G., Paterson, A.D. and Yap, A.S., 2002. Cadherin-directed actin assembly: E-cadherin physically associates with the Arp2/3 complex to direct actin assembly in nascent adhesive contacts. Curr Biol. 12, 379–82.

60. Kovacs, E.M., Verma, S., Ali, R.G., Ratheesh, A., Hamilton, N.A., Akhmanova, A. and Yap, A.S., 2011. N-WASP regulates the epithelial junctional actin cytoskeleton through a non-canonical post-nucleation pathway. Nat Cell Biol. 13, 934–43.

61. Kuhn, S. and Geyer, M., 2014. Formins as effector proteins of Rho GTPases. Small GTPases. 5, e29513.

62. Kumar, J.P., 2012. Building an ommatidium one cell at a time. Dev Dyn. 241, 136–49.

63. Lammel, U., Bechtold, M., Risse, B., Berh, D., Fleige, A., Bunse, I., Jiang, X., Klambt, C. and Bogdan, S., 2014. The Drosophila FHOD1-like formin Knittrig acts through Rok to promote stress fiber formation and directed macrophage migration during the cellular immune response. Development. 141, 1366–80.

64. Lee, H.G., Zarnescu, D.C., MacIver, B. and Thomas, C.M., 2010. The cell adhesion molecule Roughest depends on beta(Heavy)-spectrin during eye morphogenesis in Drosophila. J Cell Sci. 123, 277–85.

65. Lee, J.K., Brandin, E., Branton, D. and Goldstein, L.S., 1997. alpha-Spectrin is required for ovarian follicle monolayer integrity in Drosophila melanogaster. Development. 124, 353–62.

66. Leerberg, J.M., Gomez, G.A., Verma, S., Moussa, E.J., Wu, S.K., Priya, R., Hoffman, B.D., Grashoff, C., Schwartz, M.A. and Yap, A.S., 2014. Tension-sensitive actin assembly supports contractility at the epithelial zonula adherens. Curr Biol. 24, 1689–99.

67. Leterrier, C. and Pullarkat, P.A., 2022. Mechanical role of the submembrane spectrin scaffold in red blood cells and neurons. J Cell Sci. 135.

68. Li, S., Liu, Z.Y., Li, H., Zhou, S., Liu, J., Sun, N., Yang, K.F., Dougados, V., Mangeat, T., Belguise, K., Feng, X.Q., Liu, Y. and Wang, X., 2024. Basal actomyosin pulses expand epithelium coordinating cell flattening and tissue elongation. Nat Commun. 15, 3000.

69. Lim, D.J., 1986. Functional structure of the organ of Corti: a review. Hear Res. 22, 117–46.

70. Logan, G., Chou, W.C. and McCartney, B.M., 2022. A Diaphanous and Enabled- dependent asymmetric actin cable array repositions nuclei during Drosophila oogenesis. Development. 149.

71. Lopez-Gay, J.M., Nunley, H., Spencer, M., di Pietro, F., Guirao, B., Bosveld, F., Markova, O., Gaugue, I., Pelletier, S., Lubensky, D.K. and Bellaiche, Y., 2020. Apical stress fibers enable a scaling between cell mechanical response and area in epithelial tissue. Science. 370.

72. Lorenzo, D.N., Edwards, R.J. and Slavutsky, A.L., 2023. Spectrins: molecular organizers and targets of neurological disorders. Nat Rev Neurosci. 24, 195–212.

73. Magie, C.R., Pinto-Santini, D. and Parkhurst, S.M., 2002. Rho1 interacts with p120ctn and alpha-catenin, and regulates cadherin-based adherens junction components in Drosophila. Development. 129, 3771–82.

74. Mahajan-Miklos, S. and Cooley, L., 1994. The villin-like protein encoded by the Drosophila quail gene is required for actin bundle assembly during oogenesis. Cell. 78, 291–301.

75. Martin, A.C., Kaschube, M. and Wieschaus, E.F., 2009. Pulsed contractions of an actin-myosin network drive apical constriction. Nature. 457, 495–9.

76. Matusek, T., Djiane, A., Jankovics, F., Brunner, D., Mlodzik, M. and Mihaly, J., 2006. The Drosophila formin DAAM regulates the tracheal cuticle pattern through organizing the actin cytoskeleton. Development. 133, 957–66.

77. Matusek, T., Gombos, R., Szecsenyi, A., Sanchez-Soriano, N., Czibula, A., Pataki, C., Gedai, A., Prokop, A., Rasko, I. and Mihaly, J., 2008. Formin proteins of the DAAM subfamily play a role during axon growth. J Neurosci. 28, 13310–9.

78. Mege, R.M. and Ishiyama, N., 2017. Integration of Cadherin Adhesion and Cytoskeleton at Adherens Junctions. Cold Spring Harb Perspect Biol. 9.

79. Miao, H. and Blankenship, J.T., 2020. The pulse of morphogenesis: actomyosin dynamics and regulation in epithelia. Development. 147.

80. Millard, T.H. and Martin, P., 2008. Dynamic analysis of filopodial interactions during the zippering phase of Drosophila dorsal closure. Development. 135, 621–6.

81. Nieset, J.E., Redfield, A.R., Jin, F., Knudsen, K.A., Johnson, K.R. and Wheelock, M.J., 1997. Characterization of the interactions of alpha-catenin with alpha-actinin and beta-catenin/plakoglobin. J Cell Sci. 110 ( Pt 8), 1013–22.

82. Ozturk-Colak, A., Moussian, B., Araujo, S.J. and Casanova, J., 2016. A feedback mechanism converts individual cell features into a supracellular ECM structure in Drosophila trachea. Elife. 5.

83. Perez-Vale, K.Z. and Peifer, M., 2020. Orchestrating morphogenesis: building the body plan by cell shape changes and movements. Development. 147.

84. Pichaud, F., 2014. Transcriptional regulation of tissue organization and cell morphogenesis: the fly retina as a case study. Dev Biol. 385, 168–78.

85. Popkova, A., Stone, O.J., Chen, L., Qin, X., Liu, C., Liu, J., Belguise, K., Montell, D.J., Hahn, K.M., Rauzi, M. and Wang, X., 2020. A Cdc42-mediated supracellular network drives polarized forces and Drosophila egg chamber extension. Nat Commun. 11, 1921.

86. Pradhan, R., Kumar, S. and Mathew, R., 2023. Lateral adherens junctions mediate a supracellular actomyosin cortex in drosophila trachea. iScience. 26, 106380.

87. Qin, X., Park, B.O., Liu, J., Chen, B., Choesmel-Cadamuro, V., Belguise, K., Heo, W.D. and Wang, X., 2017. Cell-matrix adhesion and cell-cell adhesion differentially control basal myosin oscillation and Drosophila egg chamber elongation. Nat Commun. 8, 14708.

88. Quinlan, M.E., Heuser, J.E., Kerkhoff, E. and Mullins, R.D., 2005. Drosophila Spire is an actin nucleation factor. Nature. 433, 382–8.

89. Rajan, S., Kudryashov, D.S. and Reisler, E., 2023. Actin Bundles Dynamics and Architecture. Biomolecules. 13.

90. Rauzi, M., Lenne, P.F. and Lecuit, T., 2010. Planar polarized actomyosin contractile flows control epithelial junction remodelling. Nature. 468, 1110–4.

91. Ready, D.F., Hanson, T.E. and Benzer, S., 1976. Development of the Drosophila retina, a neurocrystalline lattice. Dev Biol. 53, 217–40.

92. Riedl, J., Crevenna, A.H., Kessenbrock, K., Yu, J.H., Neukirchen, D., Bista, M., Bradke, F., Jenne, D., Holak, T.A., Werb, Z., Sixt, M. and Wedlich-Soldner, R., 2008. Lifeact: a versatile marker to visualize F-actin. Nat Methods. 5, 605–7.

93. Rodriguez-Diaz, A., Toyama, Y., Abravanel, D.L., Wiemann, J.M., Wells, A.R., Tulu, U.S., Edwards, G.S. and Kiehart, D.P., 2008. Actomyosin purse strings: renewable resources that make morphogenesis robust and resilient. HFSP J. 2, 220–37.

94. Rohn, J.L., Sims, D., Liu, T., Fedorova, M., Schock, F., Dopie, J., Vartiainen, M.K., Kiger, A.A., Perrimon, N. and Baum, B., 2011. Comparative RNAi screening identifies a conserved core metazoan actinome by phenotype. J Cell Biol. 194, 789–805.

95. Roper, K., 2015. Integration of cell-cell adhesion and contractile actomyosin activity during morphogenesis. Curr Top Dev Biol. 112, 103–27.

96. Rosales-Nieves, A.E., Johndrow, J.E., Keller, L.C., Magie, C.R., Pinto-Santini, D.M. and Parkhurst, S.M., 2006. Coordination of microtubule and microfilament dynamics by Drosophila Rho1, Spire and Cappuccino. Nat Cell Biol. 8, 367–76.

97. Sahai, E. and Marshall, C.J., 2002. ROCK and Dia have opposing effects on adherens junctions downstream of Rho. Nat Cell Biol. 4, 408–15.

98. Santa-Cruz Mateos, C., Valencia-Exposito, A., Palacios, I.M. and Martin-Bermudo, M.D., 2020. Integrins regulate epithelial cell shape by controlling the architecture and mechanical properties of basal actomyosin networks. PLoS Genet. 16, e1008717.

99. Sawyer, J.K., Choi, W., Jung, K.C., He, L., Harris, N.J. and Peifer, M., 2011. A contractile actomyosin network linked to adherens junctions by Canoe/afadin helps drive convergent extension. Mol Biol Cell. 22, 2491–508.

100. Schindelin, J., Arganda-Carreras, I., Frise, E., Kaynig, V., Longair, M., Pietzsch, T., Preibisch, S., Rueden, C., Saalfeld, S., Schmid, B., Tinevez, J.Y., White, D.J., Hartenstein, V., Eliceiri, K., Tomancak, P. and Cardona, A., 2012. Fiji: an open-source platform for biological-image analysis. Nat Methods. 9, 676–82.

101. Scott, J.A., Shewan, A.M., den Elzen, N.R., Loureiro, J.J., Gertler, F.B. and Yap, A.S., 2006. Ena/VASP proteins can regulate distinct modes of actin organization at cadherin-adhesive contacts. Mol Biol Cell. 17, 1085–95.

102. Sekine, S., Tarama, M., Wada, H., Sami, M.M., Shibata, T. and Hayashi, S., 2024. Emergence of periodic circumferential actin cables from the anisotropic fusion of actin nanoclusters during tubulogenesis. Nat Commun. 15, 464.

103. Seppa, M.J., Johnson, R.I., Bao, S. and Cagan, R.L., 2008. Polychaetoid controls patterning by modulating adhesion in the Drosophila pupal retina. Dev Biol. 318, 1–16.

104. Sharkova, M., Chow, E., Erickson, T. and Hocking, J.C., 2023. The morphological and functional diversity of apical microvilli. J Anat. 242, 327–353.

105. Sherrard, K.M., Cetera, M. and Horne-Badovinac, S., 2021. DAAM mediates the assembly of long-lived, treadmilling stress fibers in collectively migrating epithelial cells in Drosophila. Elife. 10.

106. Sjoblom, B., Salmazo, A. and Djinovic-Carugo, K., 2008. Alpha-actinin structure and regulation. Cell Mol Life Sci. 65, 2688–701.

107. Solon, J., Kaya-Copur, A., Colombelli, J. and Brunner, D., 2009. Pulsed forces timed by a ratchet-like mechanism drive directed tissue movement during dorsal closure. Cell. 137, 1331–42.

108. Sousa, S., Cabanes, D., Archambaud, C., Colland, F., Lemichez, E., Popoff, M., Boisson-Dupuis, S., Gouin, E., Lecuit, M., Legrain, P. and Cossart, P., 2005. ARHGAP10 is necessary for alpha-catenin recruitment at adherens junctions and for Listeria invasion. Nat Cell Biol. 7, 954–60.

109. Spracklen, A.J., Fagan, T.N., Lovander, K.E. and Tootle, T.L., 2014. The pros and cons of common actin labeling tools for visualizing actin dynamics during Drosophila oogenesis. Dev Biol. 393, 209–226.

110. Squarr, A.J., Brinkmann, K., Chen, B., Steinbacher, T., Ebnet, K., Rosen, M.K. and Bogdan, S., 2016. Fat2 acts through the WAVE regulatory complex to drive collective cell migration during tissue rotation. J Cell Biol. 212, 591–603.

111. Sutherland, A. and Lesko, A., 2020. Pulsed actomyosin contractions in morphogenesis. F1000Res. 9.

112. Tanaka, H., Takasu, E., Aigaki, T., Kato, K., Hayashi, S. and Nose, A., 2004. Formin3 is required for assembly of the F-actin structure that mediates tracheal fusion in Drosophila. Dev Biol. 274, 413–25.

113. Tang, V.W. and Brieher, W.M., 2012. alpha-Actinin-4/FSGS1 is required for Arp2/3- dependent actin assembly at the adherens junction. J Cell Biol. 196, 115–30.

114. Thomas, C.M. and Kiehart, D.P., 1994. Beta heavy-spectrin has a restricted tissue and subcellular distribution during Drosophila embryogenesis. Development. 120, 2039–50.

115. Thomas, C.M. and Williams, J.A., 1999. Dynamic rearrangement of the spectrin membrane skeleton during the generation of epithelial polarity in Drosophila. J Cell Sci. 112 ( Pt 17), 2843–52.

116. Thomas, C.M., Zarnescu, D.C., Juedes, A.E., Bales, M.A., Londergan, A., Korte, C.C. and Kiehart, D.P., 1998. Drosophila betaHeavy-spectrin is essential for development and contributes to specific cell fates in the eye. Development. 125, 2125–34.

117. Tomlinson, A. and Ready, D.F., 1987. Neuronal differentiation in Drosophila ommatidium. Dev Biol. 120, 366–76.

118. Treisman, J.E., 2013. Retinal differentiation in Drosophila. Wiley Interdiscip Rev Dev Biol. 2, 545–57.

119. Waddington, C.H. and Perry, M.M., 1960. The ultra-structure of the developing eye of Drosophila. Proceedings of the Royal Society of London. 153, B, 155–178.

120. Wahlstrom, G., Lahti, V.P., Pispa, J., Roos, C. and Heino, T.I., 2004. Drosophila non-muscle alpha-actinin is localized in nurse cell actin bundles and ring canals, but is not required for fertility. Mech Dev. 121, 1377–91.

121. Winkelman, J.D., Bilancia, C.G., Peifer, M. and Kovar, D.R., 2014. Ena/VASP Enabled is a highly processive actin polymerase tailored to self-assemble parallel-bundled F-actin networks with Fascin. Proc Natl Acad Sci U S A. 111, 4121–6.

122. Wulf, E., Deboben, A., Bautz, F.A., Faulstich, H. and Wieland, T., 1979. Fluorescent phallotoxin, a tool for the visualization of cellular actin. Proc Natl Acad Sci U S A. 76, 4498–502.

123. Yu-Kemp, H.C., Kemp, J.P., Jr. and Brieher, W.M., 2017. CRMP-1 enhances EVL- mediated actin elongation to build lamellipodia and the actin cortex. J Cell Biol. 216, 2463–2479.

124. Zanet, J., Jayo, A., Plaza, S., Millard, T., Parsons, M. and Stramer, B., 2012. Fascin promotes filopodia formation independent of its role in actin bundling. J Cell Biol. 197, 477–86.

