## Supplemental figures and tables for "Generation and maintenance of apical rib-like actin fibers in epithelial support cells of the *Drosophila* eye"

**Supplemental data**

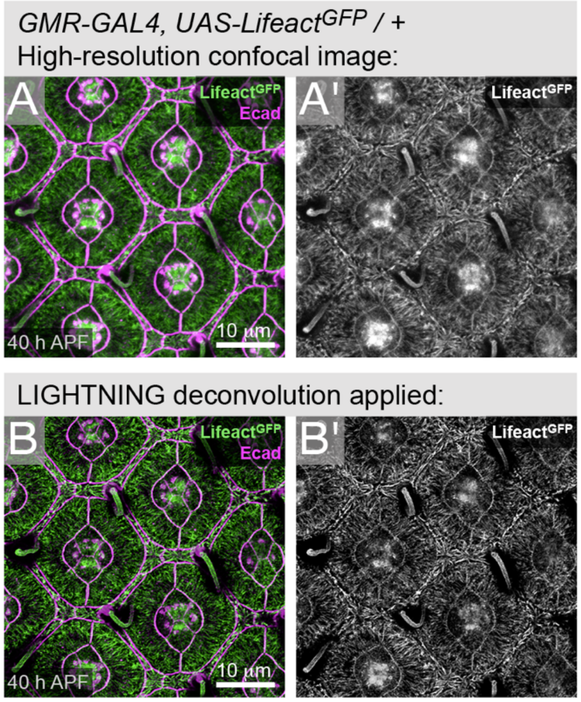


<u>Figure S1: SP8 LIGHTNING deconvolution enabled more detailed observation of the cytoskeleton in the pupal eye</u>. (A) Confocal image of a small region of an eye dissected at 40 h APF and (B) the same image after application of LIGHTNING image processing to remove signal noise and better resolve actin filaments. Scale bars are 10 $\mu$m.

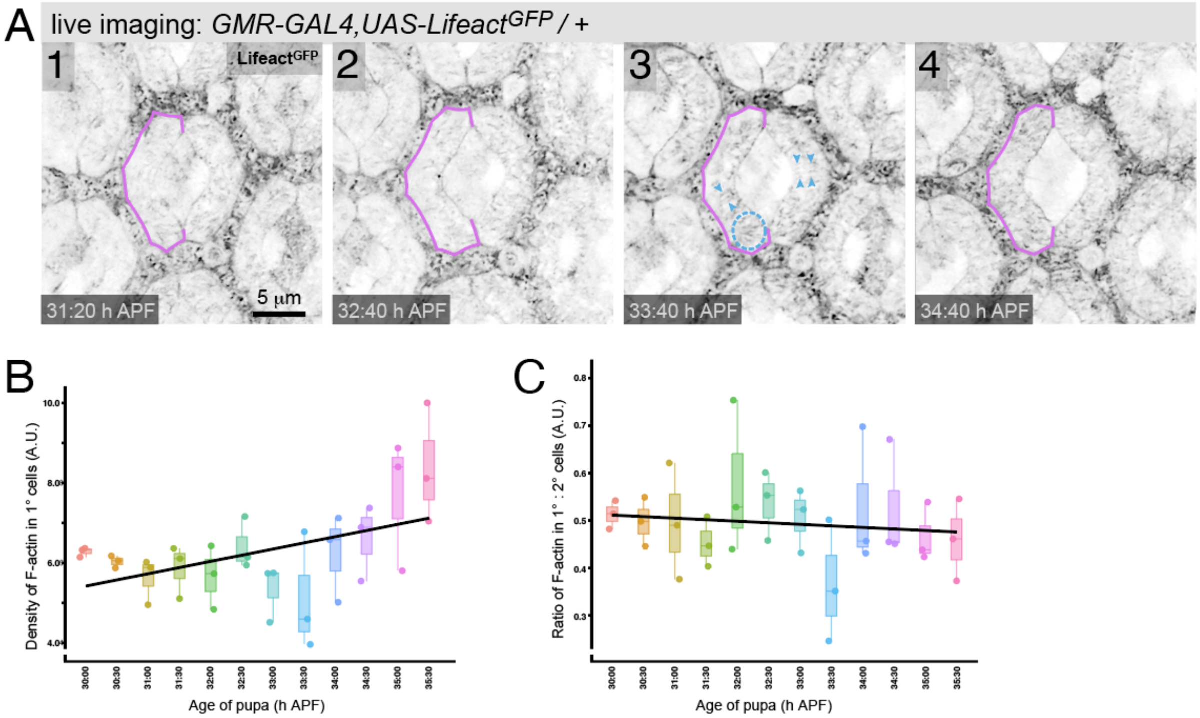


Figure S2: Filaments of the ARAF network appear from about 33 h APF. (A) Stills from a retina imaged live, with age of pupa indicated. Panel 1 and 2 are also presented in Fig. 2B. One 1° is outlined in magenta to emphasize the shape of the apical 1° cell awning. In 3, blue arrowheads indicate groups of short ARAFs oriented in parallel, and dotted circle outlines a whisker-like tuft of actin. Analyses of an eye imaged live from 30 to 35.5 h APF, as follows: (B) density of apical-cortical F-actin in 1° cells, and (C) ratio of apical-medial F-actin in 1° and 2° cells. Scale bars are 5 μm.

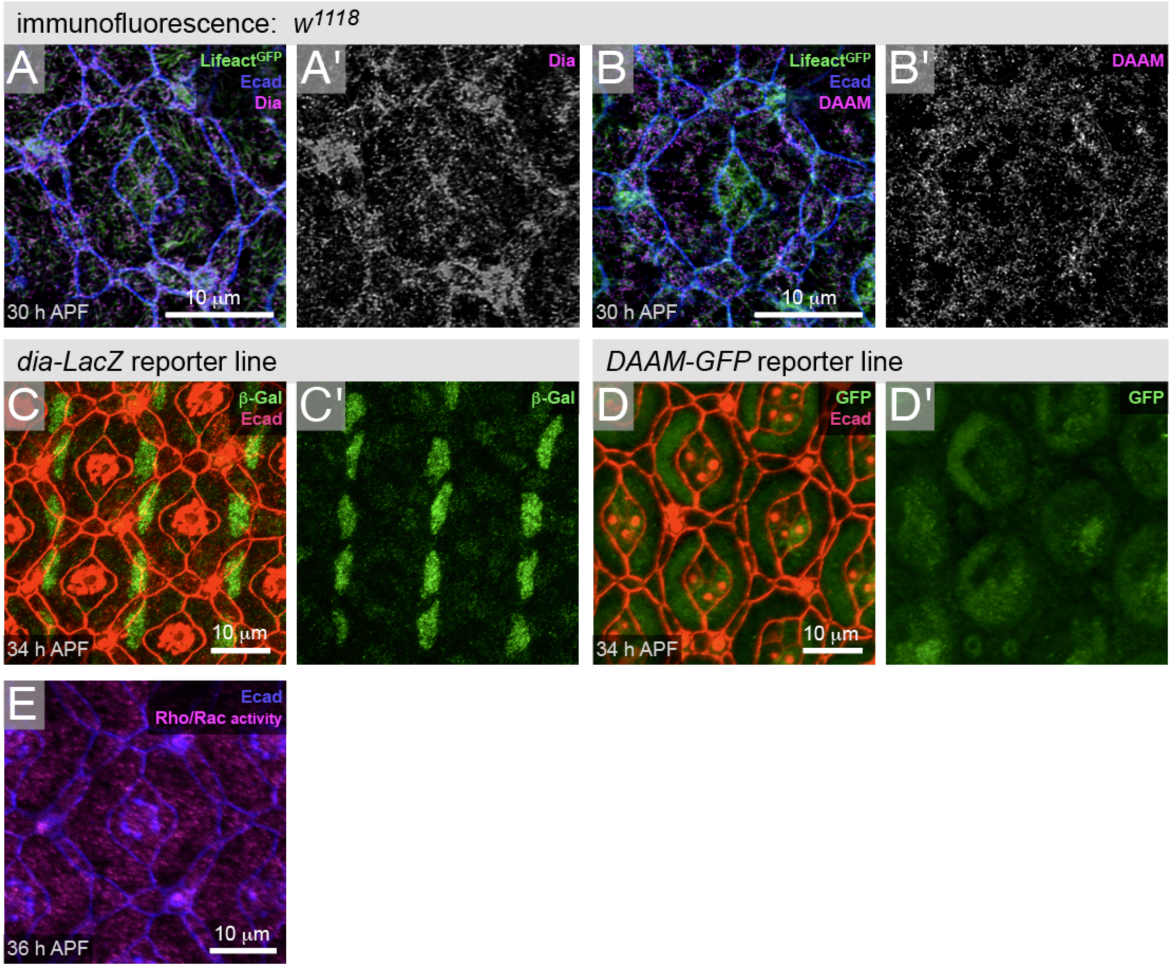


<u>Figure S3: Formins are expressed in 1° cells prior to ARAF emergence</u>. Immunofluorescence of (A) Dia and (B) DAAM at 30 h APF, before ARAFs emerge. Ecad is in blue, and Lifeact$^{GFP}$ labels the cytoskeleton. The reporter transgenes (C) *dia-lacZ* and (D) *DAAM-GFP* show robust expression of β-Gal or GFP in the nucleus (C) or cytoplasm of 1° cells (D). (E) The Rho activity sensor Pkn.RGD$^{G58A\text{-}eGFP}$, in magenta, localized through the apical region of 1° cells. Scale bars are 10 µm.

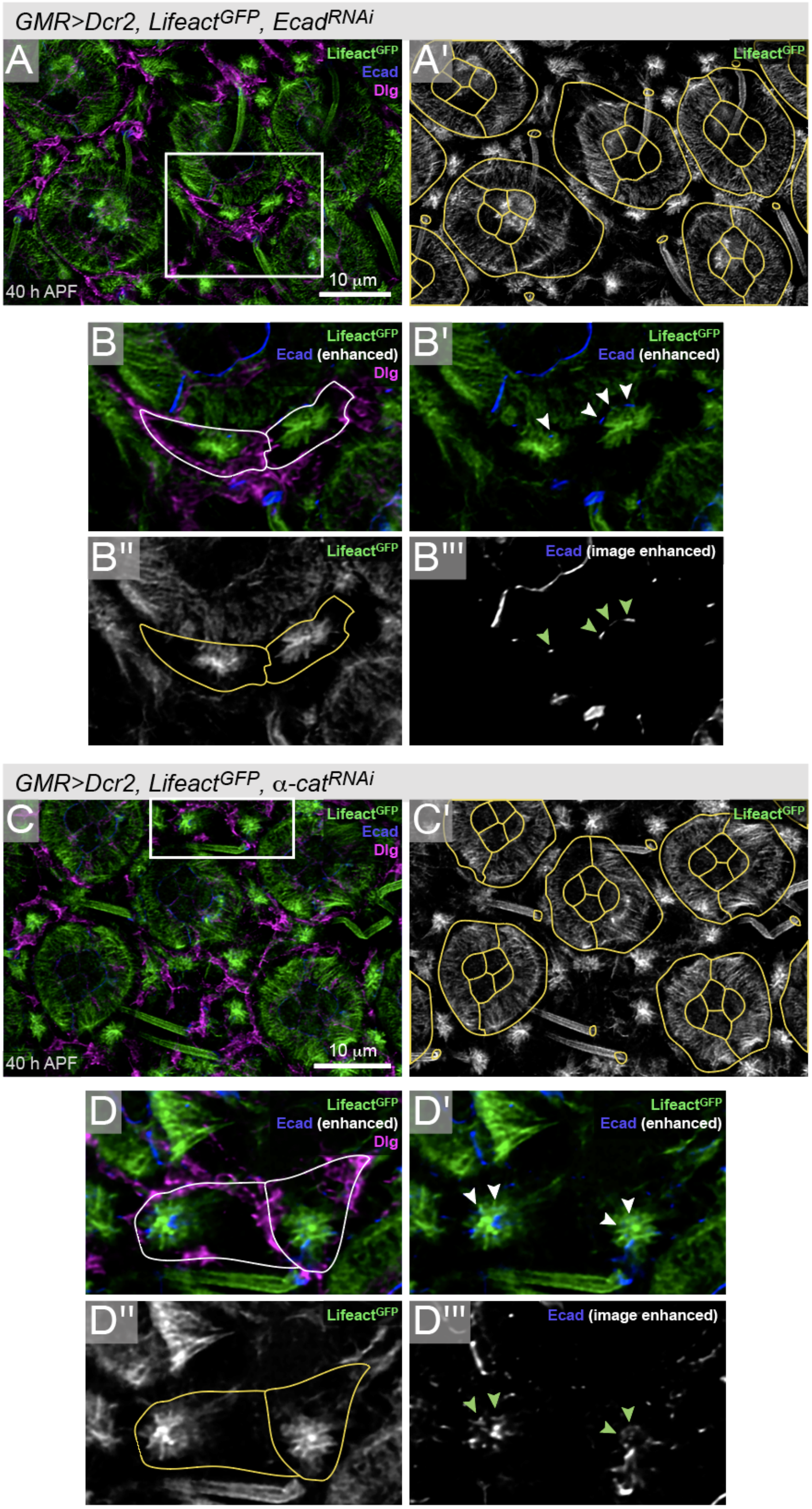
GMR>Dcr2, Lifeact^GFP, Ecad^RNAi
A
Lifeact^GFP
Ecad
Dlg
10 μm
40 h APF
A'
B
Ecad (enhanced)
B'
B''
B'''
Ecad (image enhanced)
GMR>Dcr2, Lifeact^GFP, α-cat^RNAi
C
C'
D
D'
D''
D'''

Figure S4: Integrity of AJs is essential for formation of the apical cytoskeleton in lattice cells. (A) Retina with *Ecad*$^{RNAi}$. This image is also presented in Fig. 5B. LifeactGFP labels F-actin, Ecad is in blue, and Dlg labels the lateral membrane (magenta). Ommatidia, and bristle collar cells, are traced in yellow in (A'). Boxed region is shown at higher magnification in (B). (C) Retina with a-catRNAi, with boxed region at higher magnification in (D). The outlines of two LCs are traced in white or yellow in (B) and (D) or (B'') and (D''). Ecad signal has been enhanced to visualize remaining AJs. White or green arrowheads in (B') and (D') or (B'') and (D'') indicate extant segments of AJs or internalized Ecad associated with F-actin tufts. Refer to Fig. 5(A) for control eye with *w*$^{RNAi}$ expression. Scale bars are 10 $\mu$m.

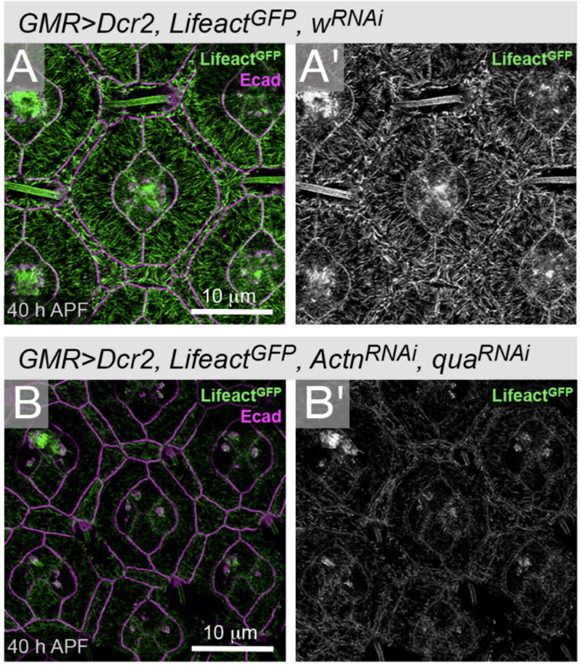


<u>Figure S5: Reducing Qua and Actn obliterates the ARAF network</u>. (A) Control *GMR>w*$^{RNAi}$ ommatidium, and (B) an ommatidia with *qua*$^{RNAi}$ and *Actn*$^{RNAi}$ expression. Image in (A) is also presented as panel (E) in Fig. 7. Scale bars are 10µm.

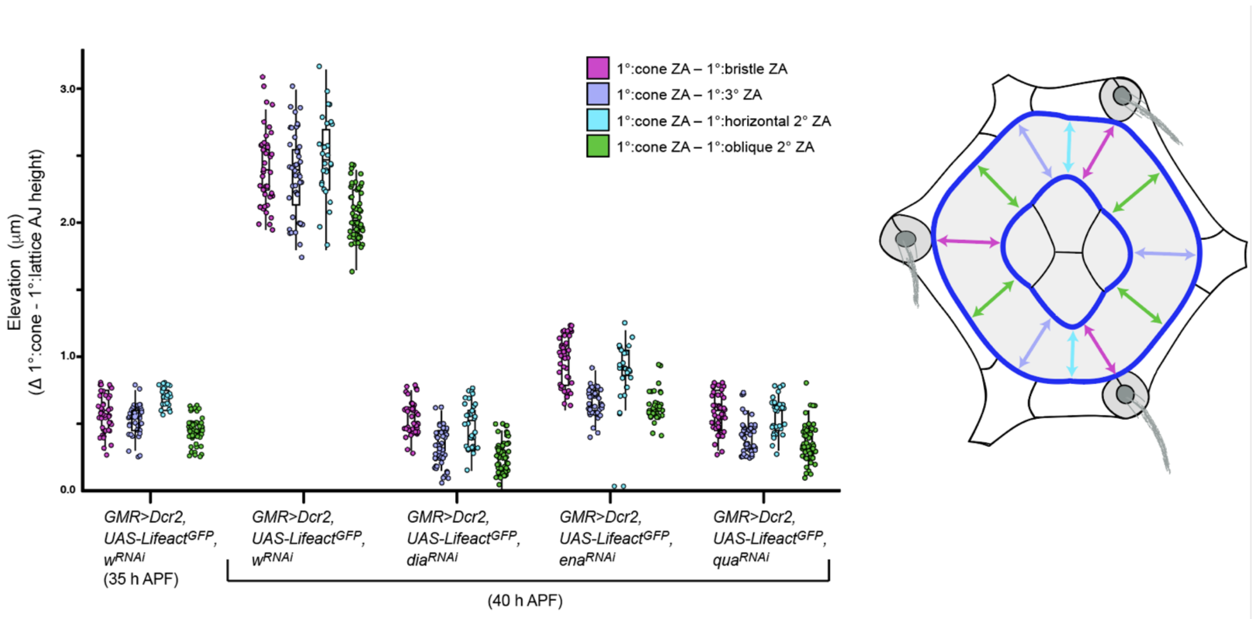


<u>Figure S6: Analyses of the elevation of the inner 1°:CC ZA, relative to the outer 1°:LC junctions</u>. These analyses separate out measurements of the change in elevation of the 1°:CC ZA, with respect to four lattice cell types, as per key and illustration at right. These data reveal that the elevation from oblique 2° cells to the 1°:CC ZA tends to be lowest and least variable.

**Supplementary Movie Titles**

Movie S1: An ommatidium in a retina imaged from 35 to 43 h APF. Lifeact$^{GFP}$ labels F-actin, driven by *GMR-GAL4* . Expression of Lifeact$^{GFP}$ was inconsistent in CCs, and bristles are also not labelled in the ommatidia shown. ARAFs become increasingly dense and organized over time. Resolution of F-actin when imaged in the live eye is lower than when imaged in tissue dissected from the pupal head.

Movie S2: Small region of an eye imaged from 45 to 55 h APF. Lifeact$^{GFP}$ labels F-actin and was not expressed in most CCs. Live-imaging data are presented as a 3-D perspective projections, with the depth-coding perspectives on the right. ARAFs become increasingly dense and ommatidial doming more pronounced through the course of this movie.

Movie S3: 3-D rendering of a pair of 1°s and the CCs of an ommatidium. 1°s (green) and CCs (pink) extend narrow projections to the base of the ommatidium that were not captured in our 3-D rendering. See also Fig.10.

Movie S4: 3-D rendering of a pair of 1°s. The 1°s connect to each other only apically, to form a collar around the CCs (not shown). See also Fig. 10.

Movie S5: 3-D rendering of a single of 1°. Each 1° curves around the CCs (not shown) connect to each other only apically to form a collar around the CCs (not shown). The shape of 1° cells is similar to that of a shark tooth. See also Fig.10.

**Table S1: Students' T-test p-values comparing angles of orientation of ARAFs (relates to Figure 1I).**

| | *ubi-Lifeact* $^{YFP}$ (n=260) | $w^{1118}$* (n=254) | *GMR>Lifeact* $^{RFP}$ (n=260) |
|---|---|---|---|
| *GMR>Lifeact* $^{GFP}$ (n=255) | 0.605 | 0.92 | 0.096 |
| *ubi-Lifeact* $^{YFP}$ | | 0.518 | 0.028 |
| $w^{1118}$* | | | 0.092 |

* ARAFs detected with rhodamine phalloidin

**Table S2: Students' T-test p-values comparing ARAF density (ARAFs/μm) (relates to Figure 1J).**

| | *ubi-Lifeact* $^{YFP}$ (n=125) | $w^{1118}$* (n=113) | *GMR>Lifeact* $^{RFP}$ (n=180) |
|---|---|---|---|
| *GMR>Lifeact* $^{GFP}$ (n=200) | $0.001x\ 10^{-12}$ | $0.008 \times 10^{-10}$ | $0.002 \times 10^{-08}$ |
| *ubi-Lifeact* $^{YFP}$ | | 0.202 | 0.071 |
| $w^{1118}$* | | | 0.566 |

* ARAFs detected with rhodamine phalloidin

**Table S3: Protein depletion in 1°cells of retinas dissected at 40 h APF.**

| Protein targetted | Transgene expressed ** | # of indepent eyes analyzed | # of 1° cells analyzed | Mean fluorescence (AU) | Standard deviation | Mean % protein reduction | p-value (t-test) |
|---|---|---|---|---|---|---|---|
| DAAM | $w^{RNAi}$ | 3 | 40 | 93.76 | 19.68 | 39.68 | $0.002 \times 10^{-18}$ |
| | $DAAM^{RNAi\text{-}GD8382}$ | 4 | 34 | 56.56 | 5.53 | | |
| Dia | $w^{RNAi}$ | 3 | 55 | 95.8 | 18.59 | 83.31 | $0.005 \times 10^{-43}$ |
| | $dia^{RNAi\text{-}KK101745}$ | 4 | 40 | 15.99 | 4.36 | | |
| Ena | $w^{RNAi}$ | 3 | 44 | 84.62 | 24.11 | 74.20 | $0.001 \times 10^{-23}$ |
| | $ena^{RNAi\text{-}GD8910}$ | 3 | 30 | 21.83 | 2.54 | | |
| Actn | $w^{RNAi}$ | 3 | 44 | 83.45 | 8.32 | 84.60 | $0.003 \times 10^{-54}$ |
| | $Actn^{RNAi\text{-}GD1354}$ | 3 | 43 | 12.85 | 3.5 | | |
| Qua | $w^{RNAi}$ | 3 | 47 | 48.01 | 4.08 | 65.42 | $0.008 \times 10^{-58}$ |
| | $qua^{RNAi\text{-}KK103765}$ | 3 | 30 | 16.6 | 2.14 | | |
| Kst | $w^{RNAi}$ | 3 | 42 | 23.86 | 3.92 | 60.79 | $0.001 \times 10^{-30}$ |
| | $kst^{RNAi\text{-}GD1472}$ | 3 | 39 | 9.35 | 1.14 | | |
| α-Spec | $w^{RNAi}$ | 3 | 41 | 56.86 | 5.08 | 29.03 | $0.008 \times 10^{-25}$ |
| | $\alpha\text{-}spec^{RNAi\text{-}HMC04371}$ | 3 | 37 | 40.31 | 5.46 | | |

**** full genotypes: *UAS-Dcr2 / + ; GMR-GAL4 / UAS-RNAi transgene***

**Table S4: Students' T-test p-values comparing LifeactGFP fluorescence intensity in 1° cells (relates to Figure 4E).**

| | *GMR>Dcr2, Lifeact $^{GFP}$, dia $^{RNAi-KK101745}$ (n=74)* | *GMR>Dcr2, Lifeact $^{GFP}$, DAAM $^{RNAi-GD8382}$ (n=42)* | *GMR>Dcr2, Lifeact $^{GFP}$, ena $^{RNAi-GD8910}$ (n=58)* |
|---|---|---|---|
| *GMR>Dcr2, Lifeact $^{GFP}$, w $^{RNAi}$ (n=60)* | $0.005 \times 10^{-48}$ | $0.007 \times 10^{-18}$ | $0.001 \times 10^{-47}$ |

**Table S5: Students' T-test p-values comparing LifeactGFP fluorescence intensity at AJs of 1° cells (relates to Figure 4F).**

| | *GMR>Dcr2, LifeactGFP, dia $^{RNAi-KK101745}$ (n=72)* | *GMR>Dcr2, Lifeact $^{GFP}$, DAAM $^{RNAi-GD8382}$ (n=72)* | *GMR>Dcr2, Lifeact $^{GFP}$, ena $^{RNAi-GD8910}$ (n=62)* |
|---|---|---|---|
| *GMR>Dcr2, Lifeact $^{GFP}$, w $^{RNAi}$ (n=71)* | $0.004 \times 10^{-34}$ | $0.003 \times 10^{-44}$ | $0.004 \times 10^{-41}$ |
| *GMR>Dcr2, LifeactGFP, dia $^{RNAi-KK101745}$* | | $0.002 \times 10^{-10}$ | $0.005 \times 10^{-08}$ |
| *GMR>Dcr2, LifeactGFP, DAAM $^{RNAi-GD8382}$* | | | 0.155 |

**Table S6: Students' T-test p-values comparing size of 1° cells (relates to Figure 4G).**

| | *GMR>Dcr2, Lifeact $^{GFP}$, dia $^{RNAi-KK10745}$ (n=74)* | *GMR>Dcr2, Lifeact $^{GFP}$, DAAM $^{RNAi-GD8382}$ (n=42)* | *GMR>Dcr2, Lifeact $^{GFP}$, ena $^{RNAi-GD8910}$ (n=58)* |
|---|---|---|---|
| *GMR>Dcr2, Lifeact $^{GFP}$, w $^{RNAi}$ (n=60)* | $0.009 \times 10^{-55}$ | $0.001 \times 10^{-43}$ | $0.001 \times 10^{-53}$ |

**Table S7: Students' T-test p-values comparing LifeactGFP fluorescence intensity in 1° cells (relates to Figure 5D).**

| | *GMR>Lifeact $^{GFP}$, $\alpha$-cat $^{RNAi-3HH}$ (n=60)* | *GMR>Lifeact $^{GFP}$, Ecad $^{RNAi[B107A1]}$ (n=57)* |
|---|---|---|
| *GMR>Lifeact $^{GFP}$, w $^{RNAi}$ (n=72)* | $0.006 \times 10^{-14}$ | $0.008 \times 10^{-25}$ |

**Table S8: Students' T-test p-values comparing size of 1° cells (relates to Figure 5E).**

| | *GMR>Lifeact $^{GFP}$, $\alpha$-cat $^{RNAi-3HH}$ (n=60)* | *GMR>Lifeact $^{GFP}$, Ecad $^{RNAi[B107A1]}$ (n=57)* |
|---|---|---|
| *GMR>Lifeact $^{GFP}$, w $^{RNAi}$ (n=51)* | $0.003 \times 10^{-10}$ | $0.003 \times 10^{-12}$ |

**Table S9: Students' T-test p-values comparing area of ARAF network (relates to Figure 5F.**

| | *GMR>Lifeact $^{GFP}$, $\alpha$-cat $^{RNAi-3HH}$ (n=25)* | *GMR>Lifeact $^{GFP}$, Ecad $^{RNAi[B107A1]}$ (n=21)* |
|---|---|---|
| *GMR>Lifeact $^{GFP}$, w $^{RNAi}$ (n=36)* | $0.002 \times 10^{-9}$ | $0.005 \times 10^{-8}$ |

**Table S10: Students' T-test p-values comparing angles between actin filaments in 1° cells (relates to Figure 5G).**

| | *GMR>Lifeact $^{GFP}$, $\alpha$-cat $^{RNAi-3HH}$ (n=131)* | *GMR>Lifeact $^{GFP}$, Ecad $^{RNAi[B107A1]}$ (n=134)* |
|---|---|---|
| *GMR>Lifeact $^{GFP}$, w $^{RNAi}$ (n=144)* | $0.004 \times 10^{-14}$ | $0.001 \times 10^{-13}$ |
| *GMR>LifeactGFP, a-cat $^{RNAi-3HH}$* | | 0.214 |

**Table S11: Students' T-test p-values comparing LifeactGFP fluorescence intensity in 1° cells (relates to Figure 6G).**

| | *GMR>Dcr2, Lifeact $^{GFP}$, $\alpha$-spec $^{RNAi\text{-}HMC04371}$* (n=42) | *GMR>Dcr2, Lifeact $^{GFP}$, kst $^{RNAi\text{-}GD1472}$* (n=70) |
|---|---|---|
| *GMR>Dcr2, Lifeact $^{GFP}$, w $^{RNAi}$* (n=96) | $0.002 \times 10^{-21}$ | $0.001 \times 10^{-37}$ |

**Table S10: Students' T-test p-values comparing size of 1° cells (relates to Figure 6H).**

| | *GMR>Dcr2, Lifeact $^{GFP}$, $\alpha$-spec $^{RNAi\text{-}HMC04371}$* (n=42) | *GMR>Dcr2, Lifeact $^{GFP}$, kst $^{RNAi\text{-}GD1472}$* (n=70) |
|---|---|---|
| *GMR>Dcr2, Lifeact $^{GFP}$, w $^{RNAi}$* (n=96) | $0.002 \times 10^{-21}$ | $0.001 \times 10^{-37}$ |

**Table S13: Students' T-test p-values comparing LifeactGFP fluorescence intensity in 1° cells (relates to Figure 7G).**

| | *GMR>Dcr2, Lifeact $^{GFP}$, Actn $^{RNAi-GD1354}$* *(n=44)* | *GMR>Dcr2, Lifeact $^{GFP}$, qua $^{RNAi-KK103765}$* *(n=46)* |
|---|---|---|
| *GMR>Dcr2, Lifeact $^{GFP}$, w $^{RNAi}$* *(n=48)* | $0.002 \times 10^{-24}$ | $0.001 \times 10^{-36}$ |

**Table S14: Students' T-test p-values comparing elevation of 1°:cone cell AJ (in relation to 1°:lattice cell AJ) (relates to Figure 9A).**

| | *GMR>Dcr2, Lifeact $^{GFP}$, w $^{RNAi}$ 40 h APF, n=168* | *GMR>Dcr2, Lifeact $^{GFP}$, dia $^{RNAi-KK101745}$ 40 h APF, n=192* | *GMR>Dcr2, Lifeact $^{GFP}$, ena $^{RNAi-GD8910}$ 40 h APF, n=153* | *GMR>Dcr2, Lifeact $^{GFP}$, qua $^{RNAi-KK103765}$ 40 h APF, n=180* |
|---|---|---|---|---|
| *GMR>Dcr2, Lifeact $^{GFP}$, w $^{RNAi}$ 35 h APF, n=180* | $0.002 \times 10^{-152}$ | $0.02 \times 10^{-24}$ | $0.001 \times 10^{-27}$ | $0.006 \times 10^{-12}$ |
| *GMR>Dcr2, Lifeact $^{GFP}$, w $^{RNAi}$ 40 h APF* | | $0.003 \times 10^{-171}$ | $0.001 \times 10^{-151}$ | $0.009 \times 10^{-165}$ |
| *GMR>Dcr2, Lifeact $^{GFP}$, dia $^{RNAi-KK101745}$ 40 h APF* | | | $0.006 \times 10^{-53}$ | $0.008 \times 10^{-2}$ |
| *GMR>Dcr2, Lifeact $^{GFP}$, ena $^{RNAi-GD8910}$* 40 h APF | | | | $0.001 \times 10^{-41}$ |

**Table S15: *Drosophila* lines utilized (excluding RNAi transgenics)**

| Abbreviated genotype | Full genotype | Source |
|---|---|---|
| *α-Spec-GFP* | *P{Wee-P.un} α-Spec[Wee-P]* | Gift from Claire Thomas, UPENN (Khanna et al., 2015) |
| *Actn-GFP* | *y[1] P{w[+mC]=PTT-GC}Actn[CC01961] w[*]/FM0, Bar[+]* | BDSC stock # 602268 |
| *coinFLP-GAL4, UAS-GFP* | *w-; P{y[+t7.7] w[+mC]=CoinFLP-GAL4}attP40 P{w[+mC]=UAS-2xEGFP}AH2* | BDSC stock # 58751 |
| *DAAM-GFP* | *y[1] w[*] Mi{PT-GFSTF.0}DAAM[MI04569-GFSTF.0] lncRNA:CR46248[MI04569-GFSTF.0-X]/FM7j, Bar[1]* | BDSC stock # 60213 |
| *dia-lacZ* | *P{ry[+t7.2]=PZ}dia[1] cn[1]/CyO; ry[506]* | BDSC stock # 11762 |
| *ey-FLP, UAS-Dcr-2 ; Sp/CyO* | *y[d2] w[*] P{ry[+t7.2]=ey-FLP.N}2 P{w[+mC]=UAS-Dcr-2.D}1; wg[Sp-1]/CyO* | BDSC stock # 58756 |
| *GMR-GAL4* | *w[*]; P{w[+mC]=GAL4-ninaE.GMR}12* | BDSC stock # 1104 |
| *GMR>Lifeact $^{GFP}$* | *w[*]; P{w[+mC]=GAL4-ninaE.GMR}12, P{y[+t*] w[+mC]=UAS-Lifeact-GFP}VIE-260B* | This study. Generated from BDSC #1104 and #35544. |
| *GMR>Lifeact $^{RFP}$* | *w[*]; P{w[+mC]=GAL4-ninaE.GMR}12, M{w[+mC]=UASp-Lifeact.TagRFP-T}ZH-22A* | This study. Generated from BDSC #1104 and #58713. |
| *kst-GFP* | *y[1] w[*]; Mi{PT-GFSTF.1}kst[MI03134-GFSTF.1]* | BDSC stock # 60193 |
| *shg $^{mTom}$* | *y[1] w[*]; TI{TI}shg[mTomato]* | BDCS stock # 58789 |
| *UAS-Dcr2* | *P{w[+mC]=UAS-Dcr-2.D}1, w[1118]* | BDSC stock # 24646 |
| *UAS-Dcr2; GMR>Lifeact $^{GFP}$* | *w[*], P{w[+mC]=UAS-Dcr-2.D}1; P{w[+mC]=GAL4-ninaE.GMR}12, P{y[+t*] w[+mC]=UAS-Lifeact-GFP}VIE-260B* | This study. Generated from BDSC #1104, #35544 and #24648. |
| *UAS-GMA* | *w[1118]; P{w[+mC]=UAS-GMA}3* | BDCS stock # 31776 |
| *UAS-Lifeact $^{GFP}$* | *y[1] w[*]; P{y[+t*] w[+mC]=UAS-Lifeact-GFP}VIE-260B* | BDSC stock # 35544 |
| *UAS-Lifeact $^{RFP}$* | *w[*]; M{w[+mC]=UASp-Lifeact.TagRFP-T}ZH-22A* | BDSC stock # 58713 |
| *ubi-Lifeact $^{YFP}$* | *P{Ubi-Lifeact.YFP}* | Gift from Maria Martin-Bermudo, CABD (Santa-Cruz Mateos et al., 2020) |
| *w $^{1118}$* | *w[1118]* | BDSC stock # 3605 |
| *sqh-Pkn.RBD $^{G58A-eGFP}$* | *w[*]; P{w[+mC]=sqh-Pkn.RBD.G58A-eGFP}312a P{sqh-Pkn.RBD.G58A-eGFP}312b* | BDSC stock # 52298 |

**Table S16: Information on transgenic RNAi lines tested and used in this study.**

| Abbreviated genotype | Full genotype | Source | Strength of knockdown in eye when driven by *GMR-GAL4* | Data presented |
|---|---|---|---|---|
| $w^{RNAi}$ | *y[1] v[1]; P{y[+t7.7] v[+t1.8]=TRiP.HMS00017}attP2* | BDSC stock # 33623 | Strong W knockdown, but no eye mis-patterning | yes |
| $\alpha$-*cat* $^{RNAi\text{-}3F}$ | *UAS-* $\alpha$*-Cat* $^{RNAi-3F}$ */ CyO* | Seppa et al., 2008 | Weak | |
| $\alpha$-*cat* $^{RNAi\text{-}3HH}$ | *UAS-* $\alpha$*-Cat* $^{RNAi-3HH}$ */ TM6b* | Seppa et al., 2008 | Strong | yes |
| $\alpha$-*spec* $^{RNAi\text{-}HMC04371}$ | *y[1] sc[*] v[1] sev[21]; P{y[+t7.7] v[+t1.8]=TRiP.HMC04371}attP40* | BDSC stock # 56932 | Strong | yes |
| $\alpha$-*spec* $^{RNAi\text{-}JF01727}$ | *y[1] v[1]; P{y[+t7.7] v[+t1.8]=TRiP.JF01727}attP2* | BDSC stock # 31209 | Weak | |
| *Actn* $^{RNAi\text{-}GD1354}$ | *w1118; P{GD1354}v7760* | VDRC stock # 7760 | Strong | yes |
| *Actn* $^{RNAi\text{-}GD1354}$ | *w1118; P{GD1354}v7762* | VDRC stock # 7762 | Weak | |
| *Actn* $^{RNAi\text{-}HMS00193}$ | *y[1] sc[*] v[1] sev[21]; P{y[+t7.7] v[+t1.8]=TRiP.HMS00193}attP2* | BDSC stock # 34874 | Weak | |
| *Actn* $^{RNAi\text{-}KK102286}$ | *P{KK102286}VIE-260B* | VDRC stock # 110719 | Weak | |
| *DAAM* $^{RNAi\text{-}GD8382}$ | *w1118; P{GD8382}v24885* | VDRC stock # 24885 | Strong | yes |
| *DAAM* $^{RNAi\text{-}HMS01978}$ | *y[1] sc[*] v[1] sev[21]; P{y[+t7.7] v[+t1.8]=TRiP.HMS01978}attP2* | BDSC stock # 39058 | Weak | |
| *DAAM* $^{RNAi\text{-}KK102786}$ | *P{KK102786}VIE-260B* | VDRC stock # 103921 | Medium | |
| *dia* $^{RNAi\text{-}KK101745}$ | *P{KK101745}VIE-260B* | VDRC stock # 103914 | Strong | yes |
| *dia* $^{RNAi\text{-}HM05027}$ | *y[1] v[1]; P{y[+t7.7] v[+t1.8]=TRiP.HM05027}attP2* | BDSC stock # 28541 | Weak | |
| *dia* $^{RNAi\text{-}HMS00308}$ | *y[1] sc[*] v[1] sev[21]; P{y[+t7.7] v[+t1.8]=TRiP.HMS00308}attP2* | BDSC stock # 33424 | Weak | |
| *dia* $^{RNAi\text{-}HMS06017}$ | *y[1] sc[*] v[1] sev[21]; P{y[+t7.7] v[+t1.8]=TRiP.HMS06017}attP40* | BDSC stock # 80437 | Weak | |
| *dia* $^{RNAi\text{-}GD9442}$ | *w1118; P{GD9442}v20518* | VDRC stock # 20518 | Weak | |
| *Ecad* $^{RNAi\text{-}A106A1}$ | *UAS-Ecad* $^{RNAi\,[A106A1]}$ | Seppa et al., 2008 | Medium | |
| *Ecad* $^{RNAi\text{-}B107A1}$ | *UAS-Ecad* $^{RNAi\,[B107A1]}$ | Seppa et al., 2008 | Strong | yes |
| *Ecad* $^{RNAi\text{-}E206A1}$ | *UAS-Ecad* $^{RNAi\,[E206A1]}$ | Seppa et al., 2008 | Medium | |
| *ena* $^{RNAi\text{-}GD8910}$ | *w1118; P{GD8910}v43056* | VDRC stock # 43056 | Strong | yes |
| *ena* $^{RNAi\text{-}GD8910}$ | *w1118; P{GD8910}v43058/CyO* | VDRC stock # 43058 | Medium | |
| *kst* $^{RNAi\text{-}GD1472}$ | *w1118; P{GD1472}v37075* | VDRC stock # 37075 | Strong | yes |
| *kst* $^{RNAi\text{-}GD1472}$ | *w1118; P{GD1472}v37074* | VDRC stock # 37074 | Medium | |
| *qua* $^{RNAi\text{-}GD11926}$ | *w1118; P{GD11926}v27623* | VDRC stock # 27623 | Weak | |
| *qua* $^{RNAi\text{-}KK103765}$ | *P{KK103765}VIE-260B* | VDRC stock # 100856 | Strong | yes |

**Table S17: Antibodies utilized in this study**

| Primary antibody | Dilution | Source | Citation |
|---|---|---|---|
| mouse anti-α-Actn | 1 in 50 | Developmental Studies Hybridoma Bank, DSHB Cat# 2G3-3D7 RRID:AB_2721943 | Saide et al, 1989 |
| mouse anti-β-galactosidase | 1 in 200 | Developmental Studies Hybridoma Bank, DSHB Cat# 40-1a RRID:AB_528100 | Ghattas et al, 1991 |
| rabbit anti-DAAM | 1 in 1000 | Gift from Jozsef Mihaly, University of Szeged | Gazsó-Gerhát et al, 2023 |
| rabbit anti-Dia | 1 in 1000 | Gift from Steven Wasserman, University of California San Diego | Castrillon and Wasserman, 1994 |
| rat anti-Ecad | 1 in 40 | Developmental Studies Hybridoma Bank, DSHB Cat# DCAD2, RRID:AB_528120 | Oda et al, 1994 |
| mouse anti-Ena | 1 in 40 | Developmental Studies Hybridoma Bank, DSHB Cat# 5G2 RRID:AB_528220 | Bashaw et al, 2000 |
| chicken anti-GFP | 1 in 10000 | Abcam Ltd, Cat# 13970 | |
| rabbit anti-Kst | 1 in 500 | Gift from Claire Thomas, The Pennsylvania State University | Thomas and Kiehart , 1994 |
| mouse anti-Qua | 1 in 50 | Developmental Studies Hybridoma Bank, DSHB Cat# 6B9 RRID:AB_528447 | Mahajan-Miklos and Cooley, 1994 |
| mouse anti-α-Spec | 1 in 50 | Developmental Studies Hybridoma Bank, DSHB Cat# 389 RRID:AB_528473 | Syu et al, 1989 |
| Secondary antibody | | | |
| Cy™3 AffiniPure™ Goat anti-Mouse IgG (H+L) | 1 in 200 | Jackson ImmunoResearch Laboratories Inc. Cat# 115-165-166 | |
| Cy™3 AffiniPure™ Goat anti-Rabbit IgG (H+L) | 1 in 200 | Jackson ImmunoResearch Laboratories Inc. Cat# 115-165-144 | |
| Alexa Fluor® 647 AffiniPure™ Donkey anti-Rat IgG (H+L) | 1 in 200 | Jackson ImmunoResearch Laboratories Inc. Cat# 712-605-153 | |
| Alexa Fluor® 488 AffiniPure™ Donkey Anti-Chicken IgY (IgG) (H+L) | 1 in 300 | Jackson ImmunoResearch Laboratories Inc. Cat# 703-545-155 | |
